# Insights from CRISPR/Cas9-mediated gene editing of centromeric histone H3 (CENH3) in carrot (*Daucus carota* subsp. *sativus*)

**DOI:** 10.1101/2022.09.19.508489

**Authors:** Frank Dunemann, Antje Krüger, Kerstin Maier, Sabine Struckmeyer

## Abstract

The generation of haploids is one of the most powerful means to accelerate the plant breeding process. In most crop species, an efficient haploid technology is not yet available or only applicable to a limited set of genotypes. Recent results published for *Arabidopsis thaliana* and major cereal crops like maize and wheat about successful haploid induction by CRISPR/Cas9-mediated editing of the centromeric histone H3 gene (CENH3) suggest that this novel method for the production of haploid plants might also be applicable to vegetable species like carrot. Here, we report and summarize the different experimental and genetic approaches that have been focused in the past few years on CRISPR/Cas9-based editing of the carrot CENH3 gene. We also describe the discovery of a second CENH3 locus in the carrot genome, which complicates the attempts to generate and to analyse putative haploid inducer genotypes. We show that three different CRISPR/Cas9 target constructs, used alone or in combinations, could successfully target carrot CENH3. Promising mutants such as in-frame indel or in-frame deletion mutants have been found, but their successful usage as putative haploid inducer is uncertain yet. Next generation sequencing of amplicons spanning CRISPR target sites and transcript-based amplicon sequencing seemed to be appropriate methods to select promising mutants, to estimate mutation frequencies, and to allow a first prediction which gene was concerned. Another aim of this study was the simultaneous knockout and complementation of the endogenous carrot CENH3 gene by an alien CENH3 gene. Co-transformation of a CRISPR/Cas9-based carrot CENH3 knockout construct together with a CENH3 gene cloned from ginseng (*Panax ginseng*) was performed by using *Rhizobium rhizogenes*. It was shown, that ginseng CENH3 protein is accumulated inside the kinetochore region of carrot chromosomes, indicating that *PgCENH3* might be a suited candidate for this approach. However, presently it is unclear, if this gene is fully functioning during the meiotic cell divisions and able to complement lethal gametes. Challenges and future prospects to develop a CENH3-based HI system for carrot are discussed.

## Introduction

The cultivated carrot (*Daucus carota* subsp. *sativus*) is one of the most widely grown vegetable crops in the world due to its aromatic flavor and high contents of health-promoting compounds including carotinoids, anthocyanins, polyacetylenes and volatile terpenes. Due to the abundant availability, high diversity of cultivars, world-wide experience and its high agricultural yields, carrot is a very promising target vegetable for attempts to enhance the efficiency of F_1_ hybrid breeding systems. Traditional inbred line production in carrots is long-lasting, and the application of tissue culture techniques such as microspore culture is highly genotype-dependent and therefore mostly inefficient. Uniparental genome elimination by manipulating centromeric histone H3 (CENH3) might be an alternative future strategy to generate fully homozygous lines for hybrid breeding. CENH3 is a protein with a universal function in chromosomes of plants and animals. As a main component of the kinetochore, a multi-protein complex to which spindle microtubules are attached during mitosis and meiosis, is involved in controlling proper segregation of chromosomes during cell division (Jiang et al. 2003). Modifications of CENH3 as one of the structural key components of centromeres could negatively affect their function and cause disturbed chromosome distribution due to loss of intact CENH3 chromatin. Manipulating CENH3 gene transcription, translation or CENH3 protein loading onto the centromeric regions of the chromosomes have been proposed as a biotechnological tool for producing haploid- and double-haploid plants (Ravi and Chan 2010; Ravi et al. 2014; Karimi-Ashtiyani et al. 2015). Therefore, CENH3-based uniparental genome elimination has become a new target for research on innovative plant breeding strategies in the past few years.

Presently, CENH3-mediated haploid induction protocols leading to a sufficiently high number of haploids have been developed solely for the model plant *A. thaliana* (Ravi and Chan 2010; Kuppu et al. 2015; Maheshwari et al. 2015; Karimi-Ashtiyani et al. 2015; Kuppu et al. 2020). However, outside of the *Arabidopsis* model system, a workable CENH3-based haploid technology has rarely been reported. An overwiew about CENH3 modifications used for haploid induction (HI) in different crop plants and the generally low achieved HI rates achieved is given by Kalinowska et al. (2019). For instance, in maize a HI rate of only 3.6% haploids was reported by Kelliher et al. (2016). Recently, higher frequencies of haploids of up to 9% were achieved in maize by using a simplified HI system based on crossing plants which were heterozygous for a CENH3 null mutation (Wang et al. 2021). In wheat, a commercially operable HI system based on paternal lines with heterozygous CENH3 frameshift mutations created by genome editing was able to produce up to 7% HI rate (Lv et al. 2020). However, all HI frequencies reported by now for crop plants are lower than those reported for *Arabidopsis* (up to > 30%).

Two different approaches have initially been proposed for CENH3-based uniparental genome elimination (Fig. 1). The first approach is based on the so called Two-Step strategy presented for the first time by Ravi and Chan (2010), where lethal *A. thaliana* knockout CENH3 mutants were successfully rescued by transformation with modified CENH3 transgenes. Later it was demonstrated by the same group that functional complementation of *A. thaliana* CENH3 null mutants and haploid induction is also possible with untagged natural CENH3 variants from progressively distant relatives such as different *Brassica* species (Maheshwari et al. 2015). The Two-Step strategy (denoted in this paper 2-STEP) would currently most likely require the use of genetic modifications and would result in transgenic haploid inducer plants. Although it is expected that the next generation would be most widely non-transgenic due to the loss of the chromosomes of the transgenic haploid-inducer plant, there is some concern about the utilization of transgenic HI parents within breeding programs. The alternative One-Step (1-STEP) approach is based on more or less targeted genetic modifications of the endogenous CENH3 gene. By creating point mutations by TILLING, which were not lethal but responsible for reduced CENH3 centromere loading and/or the generation of so-called ‘weak centromeres’, haploid genotypes were induced in barley, sugar beet and *A. thaliana* (Karimi-Ashtiyani et al. 2015; Kuppu et al. 2015). Alternatively, to the TILLING technique, genome editing techniques such as CRISPR/Cas might be used to create targeted DNA double-strand breaks (DSBs) and to knock-out and/or generate partially functional alleles of CENH3. CRISPR-based techniques have been used successfully to edit genes of interest in a large variety of model and crop plant species (Scheben et al. 2017; Li et al. 2020; Puchta et al. 2022). Due to its relatively low costs, easy design and ability of not only introducing alterations via NHEJ (Non Homologous End Joining) but also allowing gene replacements via homologues recombination (Schiml et al. 2014), CRISPR/Cas-based gene editing offers a suitable alternative to the classical mutagenesis and gene disruption approaches for functional genomics. Currently this method is based mainly on CRISPR/Cas constructs which are used for *Agrobacterium/Rhizobium*-mediated or direct gene transfer. Besides conventional genetic transformation using RNA-guided Cas endonuclease expression units, the recent establishment of using preassembled complexes of purified Cas9 protein and target gene-specific guide RNAs (Woo et al. 2015) opens up the opportunity to implement a particularly promising 1-STEP modification of CENH3 without any transgene integration (Kalinowska et al. 2019).

**Figure 1.**
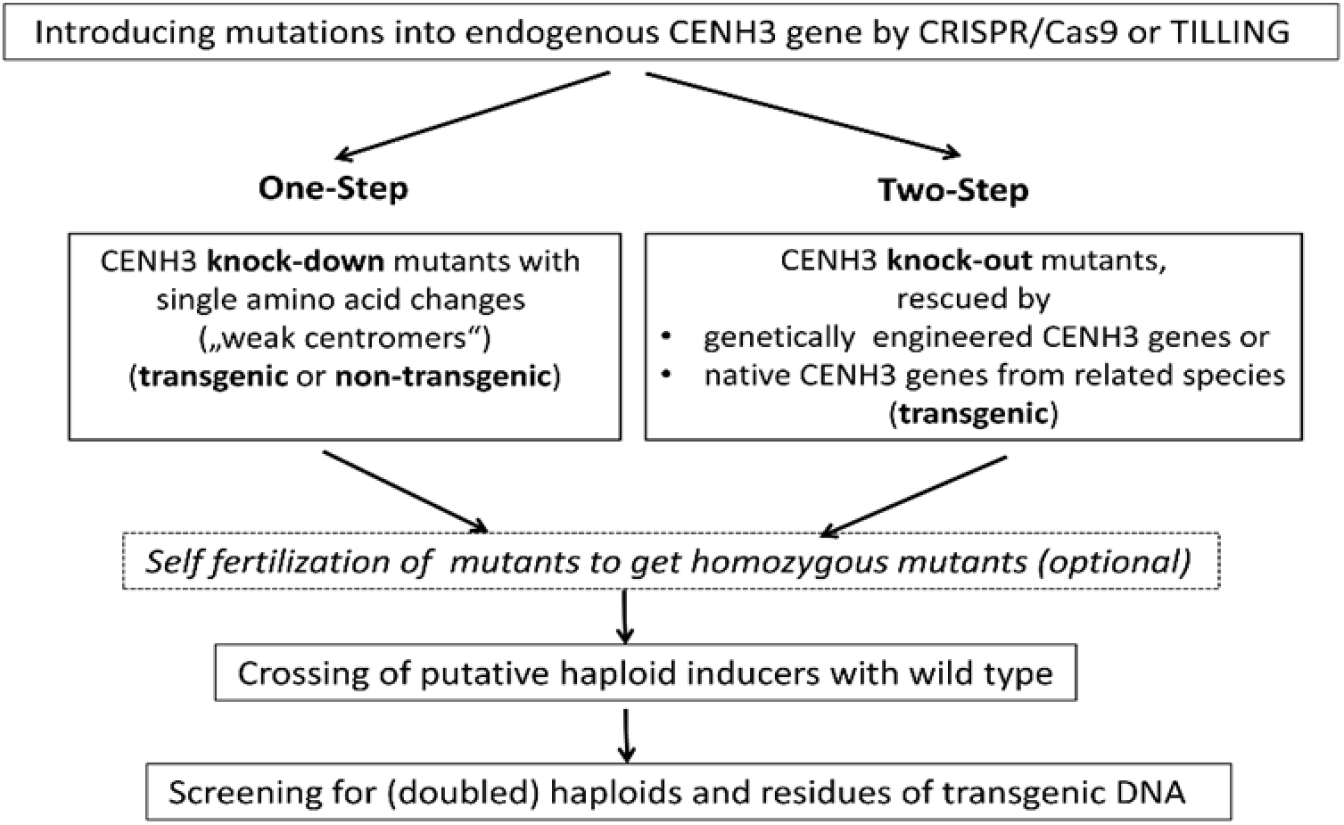
Schematic presentation of the two different strategies to create ‘haploid inducer’ genotypes through manipulation of CENH3

Successful approaches to generate haploid crop plants through CRISPR/Cas9-mediated gene editing has been reported only for the major cereal crops maize and wheat (Kelliher et al. 2019; Lv et al. 2020; Wang et al. 2021). In *A. thaliana*, CRISPR/Cas9-mediated in-frame deletions in the CENH3 histone fold domain resulted in plants with normal growth and fertility while acting as excellent haploid inducers when pollinated by wild-type pollen (Kuppu et al. 2020). In vegetable species, few studies have applied CRISPR/Cas9 technique to induce mutations in the CENH3 gene. In red cabbage (*Brassica oleracea* var. *capitata*) several sgRNAs targeting CENH3 were introduced by transfection in protoplasts and in leaves through infiltration with *Agrobacterium tumefaciens*, and average mutation rates of up to 14.4% were obtained after deep sequencing analysis (Stajič et al. 2019). Since the C-to-T mutation of the Leu133 codon (CTT) of CENH3 gene can lead to an amino acid change that resulted in haploid plants in barley, sugar beet and *Arabidopsis* (Karimi-Ashtiyani et al. 2015), this codon of the CENH3 gene of cauliflower (*Brassica oleracea* var. *botrytis*) was selected for targeted base editing. Base-edited point mutations were achieved and could be passed onto the T_1_ generation (Wang et al. 2022).

As a prerequisite for usage of a CENH3-based haploidization strategy, functional CENH3 genes of the concerned species must be cloned and analysed for the identification of suitable modification sites within the hypervariable N-terminal region of CENH3 or the more conserved histone fold domain (Kalinowska et al. 2019). In addition, for usage of the 2-STEP method, genetically engineered CENH3 genes or alien CENH3 genes have to be stably expressed in transgenic cells to complement knockout mutations of the endogenous CENH3. In carrots, functional CENH3 genes from *D. carota* subsp. *sativus* and some related *Daucus* wild species were cloned and functionally analysed (Dunemann et al. 2014). In addition, the native *PgCENH3* gene from the distantly related Apiales *s*pecies *Panax ginseng* was cloned and used for *Rhizobium rhizogenes*-based co-transformation simultaneously with a CRISPR/Cas9 construct targeting *DcCENH3* (Dunemann et al. 2019). CRISPR/Cas9-based mutations within the carrot CENH3 gene have been achieved and have been cytogenetically studied in transgenic carrot hairy root cultures and preliminary in subsequently regenerated plants (Dunemann et al. 2019). Here, we report the continuation of these studies and summarize the outcome of the 1-STEP and 2-STEP approaches after several genetic studies including self-fertilization of mutants and crossing experiments. We also describe the discovery of a second CENH3 locus in the carrot genome and report about new CENH3 knockout experiments based on additional CENH3 target sites and a second transformation of genetically homogenous transgenic carrot plants expressing already the ginseng CENH3 gene.

## Methods

### CRISPR/Cas9 constructs for targeted mutagenesis of carrot CENH3

Three target sites (called C2, C3, and C4) were selected by screening the carrot CENH3 coding region for putative target sites including potential PAM (Protospacer Adjacent Motif) sites and ruling out potential off-target binding sites using BLAST analysis (Fig. 2). Target C2 is located in exon 4 of the CDS, and target C4 is located in exon 5, respectively. The PAM of target C3 is also located in exon 5, but the target site sequence is spanning the splice site. Constructs were assembled based on the pDE-Cas9 vector backbone (Fauser et al. 2014) which provides a *bar* resistance cassette for selection. Wild-type *Rhizobium rhizogenes* strain 15834 (Lippincott et al. 1973) was transformed with the pDE-Cas9 plasmids using a modified freeze-thaw protocol according to Jyothishwaran et al. (2007). Transformed bacteria were checked by PCR and kept on CPY-S1 plates with appropriate antibiotics (for culture media composition, see Suppl. Table 1) with appropriate antibiotics as well as deep-frozen glycerol cultures for long time storage.

**Figure 2.**
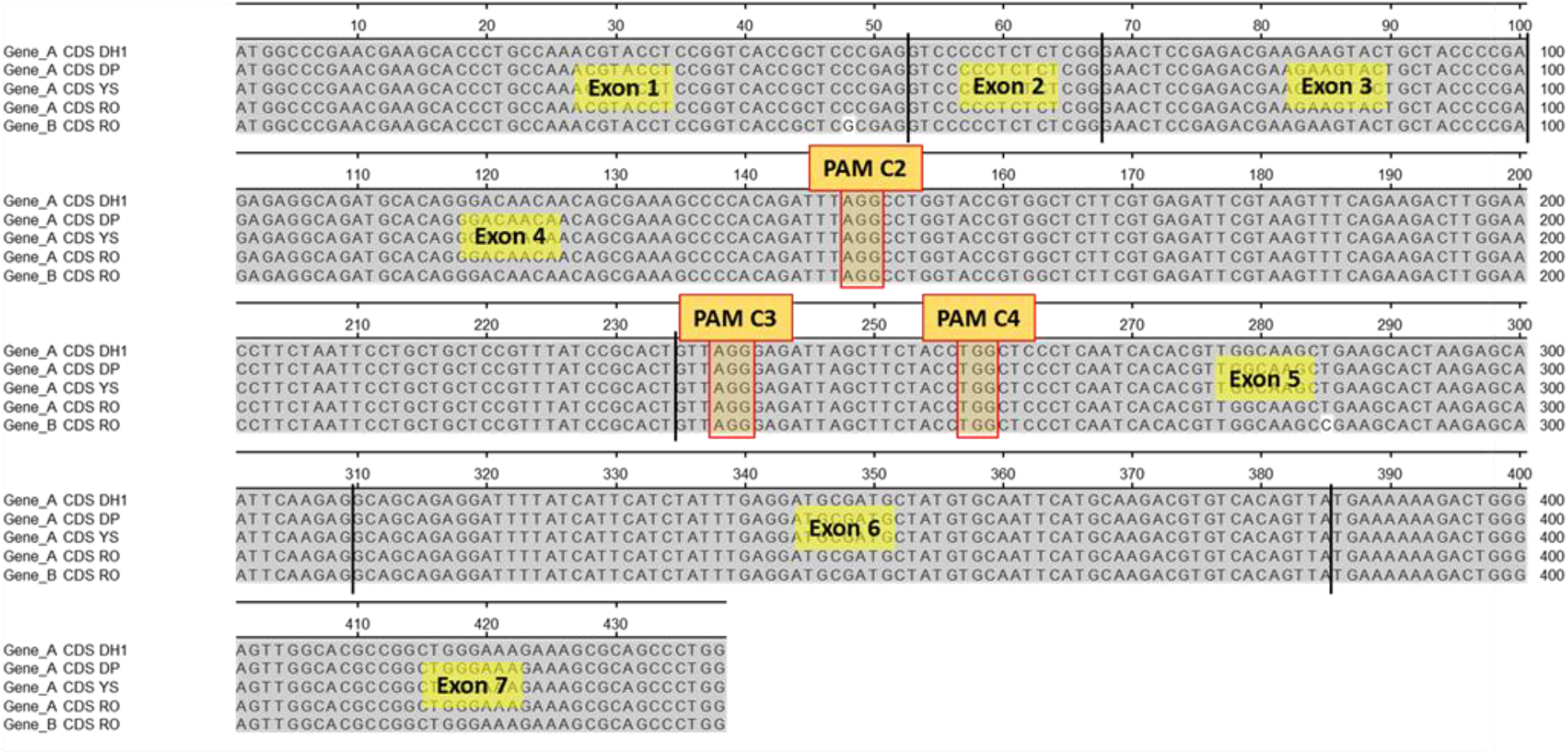
Location of CRISPR targets within the coding sequences of CENH3 of cultivars ‘Deep Purple’ (DP), ‘Yellowstone’ (YS), ‘Rotin’ (RO), and ‘DH1’ genotype used by Iorizzo et al. (2016) for carrot genome sequencing

### Ginseng CENH3 construct

To identify CENH3 orthologous sequences of *Panax ginseng*, the assembled *P. ginseng* transcriptome (Li et al. 2013) was used for *in silico* gene mining. A single contig was found spanning the whole putative CDS of *PgCENH3*. For cloning of *PgCENH3*, a PCR primer pair was designed from the 5’-UTR and 3’-UTR sequences of the contig, respectively, to amplify the whole CDS. DNA fragments with the expected size obtained after PCR with *P. ginseng* cDNA templates were blunt-end cloned by the CloneJET PCR cloning kit (ThermoFisher Scientific). The construction of the binary expression vector p6i-d35S∷*PgCENH3* was described by Dunemann et al. (2019).

### Transformation of carrot with *Rhizobium rhizogenes*

Carrot (*D. carota* subsp. *sativus*) cultivars ‘Deep Purple’ (DP), ‘Yellowstone’ (YS), ‘Rotin’ (RO) and ‘Blanche’ (BL) were used for the 1-STEP- and 2-STEP-experiments of this study. Plants obtained from seeds were grown in pots in a greenhouse until they were used in transformation experiments. In addition, transgenic T_1_ carrot plants obtained from the 2-STEP experiment were used for a second transformation with CRISPR construct combinations. Tap roots were washed, peeled and surface-sterilized by immersion in 70% ethanol for 5 min followed by a 20 min treatment with 4% sodium hypochlorite and 0.1% Tween 20. Each carrot was sliced into several 0.5 – 0.8 cm thick discs and the dead tissue of the outer edge of the discs was removed. The apical (shoot-averted) side was labeled with a sterile toothpick, and the carrot discs were placed (apical side up) onto 2% water agar (Duchefa, Haarlem, Netherlands) without any antibiotics. Bacterial inocula were prepared from liquid cultures grown in CPY-S2 medium (Suppl. Table 1) for about 24 h at 28°C in the dark on a rotary shaker (180 rpm). Cultures with an optical density (OD_600_) of 0.6 - 0.8 were either used as single inoculum in the 1-STEP experiment or mixed 1:1 for the co-transformation experiments. Table 1 summarizes the different transformation experiments and the binary plasmid combinations used for co-transformations. The transformation experiment using selected *PgCENH3*-transgenic carrots for a second transformation with combinations of two CRISPR targets is called in this study 2-STEP-DT (double transformation). About 100 μl of the inoculum was spread along the cambial ring on the upper surface of the root discs. After co-cultivation for seven days the carrot discs were placed for up to five weeks on 2% water agar supplemented with 200 mg/l cefotaxime and 100 mg/l carbenicillin (WA-CC medium, Suppl. Table 1).

**Table 1.** Transformation experiments

| Approach | Co-transformation | Binary plasmids |
| --- | --- | --- |
| 1-STEP | no | pDE-Cas9::C2 or<br>pDE-Cas9::C3 or<br>pDE-Cas9::C4 |
| 2-STEP | yes | pDE-Cas9::C4 + p6i-d35S::PgCENH3 |
| 2-STEP-DT | yes | pDE-Cas9::C2 + pDE-Cas9::C3 or<br>pDE-Cas9::C2 + pDE-Cas9::C4 |

Hairy roots started to regenerate generally after two weeks and were harvested weekly alongside with a small piece of original root tissue. Regenerated hairy roots were placed in small petridishes (6 cm diameter) on selective MS-CCP medium with 10 mg/l phosphinotricin (1-STEP and 2-STEP-DT) and cultured in the dark at 22°C. For 2-STEP co-transformation, roots were placed on selection medium with 50 mg/l hygromycin (for *PgCENH3* transgene selection) and 10 mg/l phosphinotricin (for CRISPR target selection). Hairy root cultures developing from a single excised root were considered to have an independent origin of regeneration. They were placed in a 6-cm single petridish and were designated as ‘hairy-root line’. About four weeks later the surviving and evidently propagating hairy root lines were subcultured on the same selective culture medium. At least three successive subcultures grew on the same culture medium. Then the cultures were considered to be free of remaining *R. rhizogenes* bacteria and subcultured on medium without cefotaxime and carbenicillin. PCR tests for *R. rhizogenes* contamination were performed for selected lines with PCR primer for the *virD2* gene, which is located on the *Ri* plasmid outside of the T-DNA.

### Plant regeneration from carrot hairy roots

Hairy root lines propagated over a period of about 3-4 months were used as starting material for plant regeneration via somatic embryogenesis. Segments of hairy roots with a length of about 1-2 cm were dissected and placed on plates with a medium containing half concentration of the Murashige and Skoog (MS) basic medium and 1 mg/l of the plant hormone 2,4-D (2,4-dichlorophenoxyacetic acid) for callus induction. Culture conditions were 16 h light (25°C) and 8 h darkness (22°C). Callus pieces were transferred after one or two culture passages on callus-induction medium to the same medium without 2,4-D but with kinetin (0.2 mg/l). After 4-5 weeks the callus (partly already with small somatic embryos) was transferred to hormone-free MS medium. Regenerated plantlets were sub-cultured in 10 cm glass vessels on the same medium and then transferred to soil for further growth in a greenhouse cabin. Individual plants regenerated from different regions of the embryogenic callus cultures were labeled with an individual plant number.

### Molecular analyses

Total genomic DNA of hairy roots or young leaf tissue of individual regenerated plants was extracted with the innuPREP Plant DNA kit (Analytik Jena, Jena, Germany) using 50-100 mg fresh tissue as starting material for tissue disruption by a swing mill. In order to assess the genetic status of the hairy roots and putatively transformed plants, different PCR primer were applied. To check the integration of the CRISPR/Cas9 expression cassette containing the sgRNA targeting the C2, C3 or C4 region of *DcCENH3*, the primer pair SS42/43 (Fauser et al. 2014) was used resulting in a fragment of 1070 bp in positive transformants. Additionally, PCR analysis with the primer pair BAR-F/R amplifying a 500 bp long fragment from the *bar* resistance cassette of pDe-CAS9 plasmid was performed (for PCR primer details, see Suppl. Table 2). To verify the integration of *PgCENH3* in hairy roots and regenerated plants, a primer pair was designed based on the *PgCENH3* cDNA sequence, which amplifies a 300 bp fragment in transgenic hairy roots and plants. This primer pair (PCEN-F/-R) is specific for expressed ginseng CENH3 and could not amplify the native carrot CENH3 gene. Standard PCR analysis was generally carried out in a total volume of 25 μl containing 40 ng template DNA, 1 U of ‘DreamTaq’ DNA polymerase (Thermo Fisher Scientific), 1x *Taq* polymerase buffer with 20 mM MgCl_2_ (Thermo Fisher Scientific), 0.2 μM of each primer and 0.2 mM of each dNTP. Amplification conditions were as followed: 1 cycle of 3 min at 94°C; 35 cycles of 94°C for 30 sec, 55°C-60°C for 45 sec, 72°C for 1 min; final extension of 72°C for 5 min. Specific annealing temperatures of the used primers are shown in Suppl. Table 2. PCR fragments were separated on standard 1% agarose gels at 100 V and stained using ethidium bromide.

For mutation analysis by next generation amplicon sequencing of the C2 target region, the primer pair C2-AEZ-F/-R was used, which amplifies a 400 bp fragment (Suppl. Table 2). For amplicon sequencing of the C3/C4 target region, the primer pair C4-AEZ-F/-R was used for amplification of a 369 bp fragment spanning both target sites. To allow amplicon sequencing of up to ten pooled amplicons, six nt’s were added at the 5’ end of the forward primer to form a barcode-tag (Suppl. Table 2). These barcodes should allow the mutant screening for up to ten plants in a single sequencing reaction. PCR for each target site analysis was performed in a total volume of 100 μl (2 x 50 μl). The PCR reactions were loaded onto a 1% agarose gel, and obtained fragments were cut from the gel and purified with the innuPREP Gel Extraction kit (Analytik Jena, Jena, Germany). Extracted DNA was finally diluted in 30 μl extraction buffer and quantified by a Qubit fluorometer (Thermo Fisher Scientific). Illumina-based sequencing was performed by Genewiz (Leipzig, Germany) using the service type ‘Amplicon-EZ’, which is suited for amplicons 150-500 bp in size. The option ‘variant detection’ was used for first sequence analyses. Further sequence analyses were performed with BioEdit (vers.7.2.6.1) and MEGA-X (vers.10.1.7). BioEdit was used for filtering of sequences produced by a special barcode primer and elimination of all sequences showing the wild-type target sequences. After sorting and evaluation of frequency and kind of induced mutations, FASTA files were generated and used for MEGA-X-based sequence alignments by the MUSCLE algorithm.

For amplicon sequencing of CENH3 transcripts without pooling, a universal primer pair (DCEN-F/-R) was designed that amplifies the complete CDS (minus 1 nt from the stop codon) of *DcCENH3* (Suppl. Table 2). For RNA isolation, young leaves were flash frozen in liquid nitrogen and grounded to fine powder using a swing mill. Total RNA was isolated using the innuPREP Plant RNA kit (Analytik Jena, Jena, Germany). The integrity of the RNA was checked on standard 1% agarose gels at 100 V stained using EthBr, and its concentration was measured using a NanoDrop device. Qualitatively and quantitatively checked RNA was then used to synthesize cDNA with the RevertAid H minus First Strand cDNA Synthesis Kit (ThermoFisher Scientific). This cDNA (1 μl, ~20 ng) was used for PCR with the DCEN primer pair amplifying a fragment with a length of 437 bp. Sequencing and data evaluation was done as decribed above.

### Cloning of full-length genomic DNA sequences of *DcCENH3A and DcCENH3B*

A PCR-based cloning strategy was used to clone the full-length genomic sequences (~4.500 bp) of *DcCENH3B* of ‘Rotin’ and *DcCENH3A* of the DH line (double-haploid) ‘DH1’ used by Iorizzo et al. (2016) for sequencing the carrot genome. Seeds of ‘DH1’ were kindly provided by Th. Nothnagel (JKI, Quedlinburg, Germany). After several intermediate cloning steps using the pGEM-T Easy Vector System (Promega, Madison, USA), final forward primer were designed from the 5’UTR (*DcCENH3B*) or the first nt’s of the CDS (*DcCENH3A*) that were able to amplify the whole genomic sequences when used together with an identical reverse primer (CEN-UNI-R) located in the 3’ UTR. For PCR and subsequent cloning steps, the TOPO™ XL-2 Complete PCR Cloning Kit (Invitrogen) was used. Sequencing of the full-length sequences was performed by a primer-walking approach using plasmid DNA and six intermediate primer for sequencing (Microsynth SeqLab, Göttingen, Germany).

### Cytogenetic analyses by CENH3 immunostaining

To prove the existence of carrot and ginseng CENH3 in the centromeres of mitotic root or leaf cells by immunofluorescence experiments, nuclei were isolated from 10 mg (young leaves) to 100 mg (hairy roots or segments of taproots) plant material. The plant material was kept 5 min under vacuum in 10 ml ice-cold 4% formaldehyde followed by fixation on ice for 15 min. After two washing steps in 5 ml ice-cold Tris buffer Kl (10 mM Tris HCl, 10 mM Na_2_-EDTA, 100 mM NaCl, pH7.5) for 10 min, the material was chopped up with a razor blade in 1 ml ice-cold LB01 buffer (15 mM Tris-HCl, 2 mM Na_2_-EDTA, 0.5 mM Spermin x 4 HCl, 80 mM KCl, 20 mM NaCl, 0.1% Triton X-100, 15 mM mercaptothanol, pH 7.5) and filtrated through a 35 μm mash. The filtrate was centrifuged in a 1.5 ml tube at 3.000 rpm for 3 min to facilitate the formation of nuclei clumps, which allowed a faster microscopic screening for the evaluation of the respective CENH3 in a putative CENH3 mutant. After removal of the supernatant 100 μl nuclei suspension was mixed with the same volume sucrose buffer (10 mM Tris, 50 mM KCl, 2 mM MgCl_2_, 5% (m/v) sucrose, 50 μl Tween 20, pH7.5) and left to dry overnight on a microscope slide at room temperature. The immunostaining procedure based on specific polyclonal antibodies (LifeTein, Hillsborough, USA) corresponding to the N-terminus of *DcCENH3* or *PgCENH3*, respectively (Fig. 3), was performed for each genotype separately as described by Dunemann et al. (2014). A DyLight®594-conjugated goat anti-rabbit IgG antibody (Abcam, Cambridge, UK) was used as secondary antibody for staining carrot CENH3, and a CF™488A-conjugated goat anti-rabbit IgG antibody (Sigma-Aldrich) was used for the indirect staining of ginseng CENH3. Double immunostaining of the same nuclei preparation was also performed, but this method was not used as a general screening method for the evaluation of the existence and strength of carrot and ginseng CENH3 signals.

**Figure 3.**
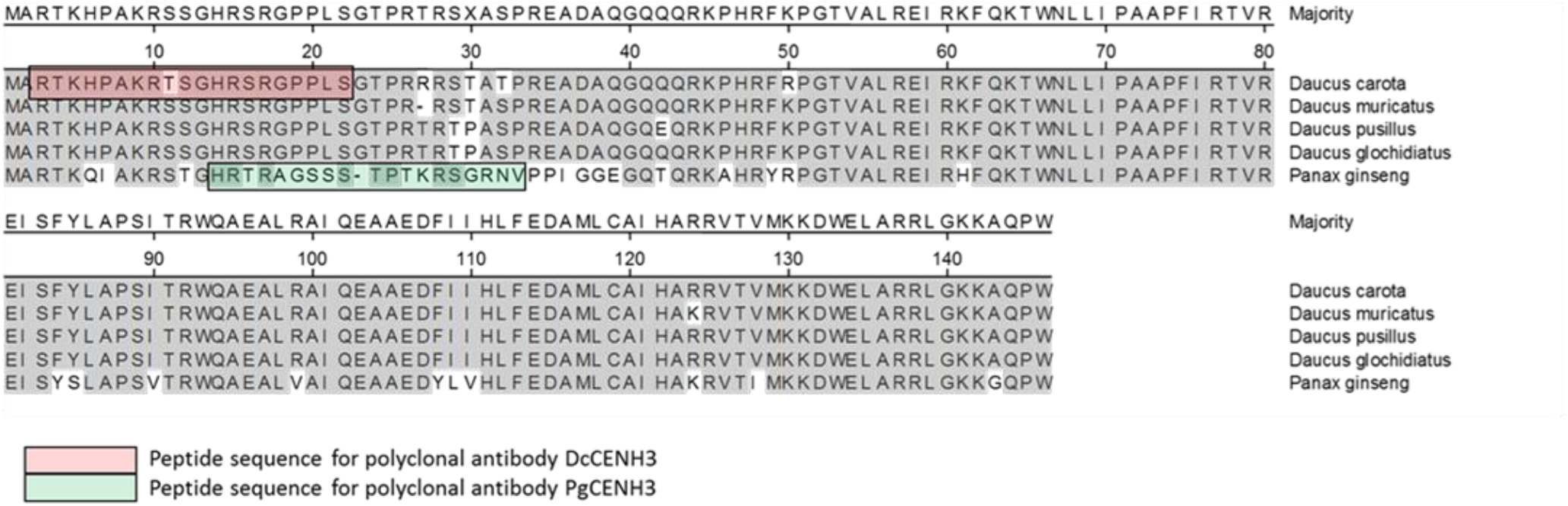
CENH3 protein sequences of *D. carota*, some other *Daucus* species, and *Panax ginseng*

## Results and discussion

Previous research on CRISPR/Cas9-based mutagenesis of carrot CENH3 has demonstrated, that CRISPR-induced mutations in the *DcCENH3* gene could be obtained with the C4 CRISPR target construct in hairy root cultures obtained after transformation with *R. rhizogenes* using both 1-STEP and 2-STEP approaches (Dunemann et al. 2019). Analysis of mutations in hairy roots and regenerated T_0_ plants indicated that mutations were mostly chimeric, which is a common phenomenon after CRISPR/Cas9-mediated mutagenesis (Schedel et al. 2017; Lee et al. 2019). After cytogenetic analyses of mutated hairy roots it was found that the CENH3 production in some hairy root lines was impaired. Although there was no relationship between the kind and extent of the mutations in the hairy roots and the CENH3 phenotype, it was obvious that mutations within the *DcCENH3* gene were associated with a reduced CENH3 protein accumulation in the centromeres of some hairy root lines (Dunemann et al. 2019). After first 2-STEP transformations, the ginseng *PgCENH3* gene was present and expressed in almost all hairy roots indicating that co-transformations have been induced efficiently in carrots. Among the first regenerated plants, no plant was found with a complete failure of carrot CENH3 signals, neither in the 2-STEP experiments (where such a finding might be possible due to the expected “rescue”-effect by ginseng CENH3) nor in the 1-STEP plants (which is expected, since complete *DcCENH3* knockout individuals would be not able to survive). To promote the generation of homogeneously mutated plants transformed with the 1-STEP or the 2-STEP constructs, T_0_ plants were crossed with several carrot cultivars and breeding lines. Although the T_0_ parents used for crossing experiments were most probably still heterozygous and/or chimeric for mutations within the carrot CENH3 gene, the generation of haploid genotypes could not be completely ruled out. Recently, it was shown that maize lines which were heterozygous for a CENH3 null mutation (containing a premature stop codon) were able to induce up to 9% haploids in progenies obtained after crosses with wild-type plants (Wang et al. 2021). However, the analysis of carrot T_1_ plants derived from crosses with T_0_ plants containing lethal CENH3 mutations did not result in haploid plants. In the following, we report about the continuation and extension of previous work on 1-STEP and 2-STEP approaches.

### 1-STEP experiments

The proven *R. rhizogenes-*based transformation system and the pDE-Cas9 vector system (Fauser et al. 2014) was used to transform different carrot cultivars with three different sgRNAs to target specific regions of the *DcCENH3* gene. The targeted region of the previously used vector pDE-Cas9∷C4 is completely on exon 5, whereas the newly used targets are either on exon 4 (pDE-Cas9∷C2) or are targeting a putative splicing site (pDE-Cas9∷C3) (Fig. 2). Based on pre-tested mutated hairy root lines, totally 120 T_0_ plants were regenerated *via* somatic embyogenesis. Out of the transformation with the C2 construct, 20 plants were obtained from originally four selected hairy root lines. The transformation with the C3 target vector resulted in 35 T_0_ plants regenerated from five hairy root lines. Including the plants derived from the initial experiment, totally 65 T_0_ plants were obtained for target C4. Almost all T_0_ plants have been sequenced by next generation amplicon sequencing using a pooling strategy with specific 6-nt barcodes for each individual plant. By this way, we have been able to analyse the sequences around the CENH3 target sites for up to ten plants simultaneously in a single sequencing reaction, which resulted in a drastic reduction of the sequencing costs. Nevertheless, we were able to use up to several thousands of sequences of each individual plant for mutation analyses. With the exception of eight plants regenerated from the hairy root line C3/BL-5, which were *DcCENH3* wild-type, all other sequenced T_0_ plants showed different types of mutations (Fig. 4). Beside many single base substitutions which we called “weak” mutations, also several “strong” mutations (single nt insertions or deletions leading to frame shift mutations, different longer insertions and deletions) were found, partly in relatively high frequency of up to 70%. For example, among eight T_o_ plant regenerated from the hairy root line C4/YS-23, the mutation frequency varied from 40 to 70% (with a mean of 56%; data not shown). The most frequent mutation in the C4/YS-23 regenerates was the insertion of an additional base ‘T’ at the position 4 bp upstream to the PAM of target C4 leading to a lethal frameshift mutation due to a premature stop codon. In most cases, more than one mutation was found in a single plant (up to four or sometimes even more) indicating that the original hairy roots as well as plants/tissues were strongly chimeric concerning the CRISPR effects. Several T_0_ plants appeared to carry biallelic mutations, as for example plant C4/YS-21-4, which had a high total mutation frequency of >60% and showed both mutations, insertion ‘T’ and deletion ‘T’, with a similar frequency (Table 2, Fig. 4).

**Table 2.** Amplicon sequencing results of progeny 19.032 and parental genotype C4/YS-21-4 used for self-fertilization

| Plant | Generation | Amplicon sequencing |  |  |  | No. of sequences |  | No. of sequences |  |
| --- | --- | --- | --- | --- | --- | --- | --- | --- | --- |
|  |  | Pooling: yes / no | Total no. of sequences | Mutated sequences | % | Insertion T | % | Deletion T | % |
| 19.032-1 | T<sub>1</sub> | yes | 2726 | 2555 | 93,7 | 1258 | 49,2 | 1187 | 46,4 |
| 19.032-2 | T<sub>1</sub> | yes | 5717 | 5330 | 93,2 | 3368 | 63,2 | 1712 | 32,1 |
| 19.032-2 | T<sub>1</sub> | no (repetition) | 28117 | 28047 | 99,8 | 17048 | 60,8 | 10051 | 35,8 |
| 19.032-3 | T<sub>1</sub> | yes | 3530 | 3334 | 94,4 | 3034 | 91,0 | 161 | 4,8 |
| 19.032-3 | T<sub>1</sub> | no (repetition) | 23290 | 23203 | 99,6 | 22324 | 96,2 | 25 | 0,1 |
| 19.032-4 | T<sub>1</sub> | yes | 2761 | 2519 | 91,2 | 1616 | 64,2 | 792 | 31,4 |
| 19.032-5 | T<sub>1</sub> | yes | 1252 | 1173 | 93,7 | 664 | 56,6 | 366 | 31,2 |
| 19.032-5 | T<sub>1</sub> | no (repetition) | 24293 | 24214 | 99,7 | 14998 | 61,9 | 8370 | 34,6 |
| 19.032-6 | T<sub>1</sub> | yes | 1585 | 1479 | 93,3 | 1356 | 91,7 | 56 | 3,8 |
| 19.032-6 | T<sub>1</sub> | no (repetition) | 28141 | 28043 | 99,7 | 27021 | 96,4 | 12 | 0,1 |
| 19.032-7 | T<sub>1</sub> | yes | 2007 | 1894 | 94,4 | 989 | 52,2 | 543 | 28,7 |
| 19.032-9 | T<sub>1</sub> | yes | 2507 | 2321 | 92,6 | 1233 | 53,1 | 670 | 28,9 |
| 19.032-11 | T<sub>1</sub> | yes | 5786 | 3289 | 56,8 | 2901 | 88,2 | 177 | 5,4 |
| 19.032-14 | T<sub>1</sub> | yes | 1370 | 898 | 65,5 | 455 | 50,7 | 361 | 40,2 |
| C4/YS-21-4 | T<sub>0</sub> (parent) | yes | 539 | 343 | 63,6 | 169 | 49,3 | 154 | 44,9 |

**Figure 4.**
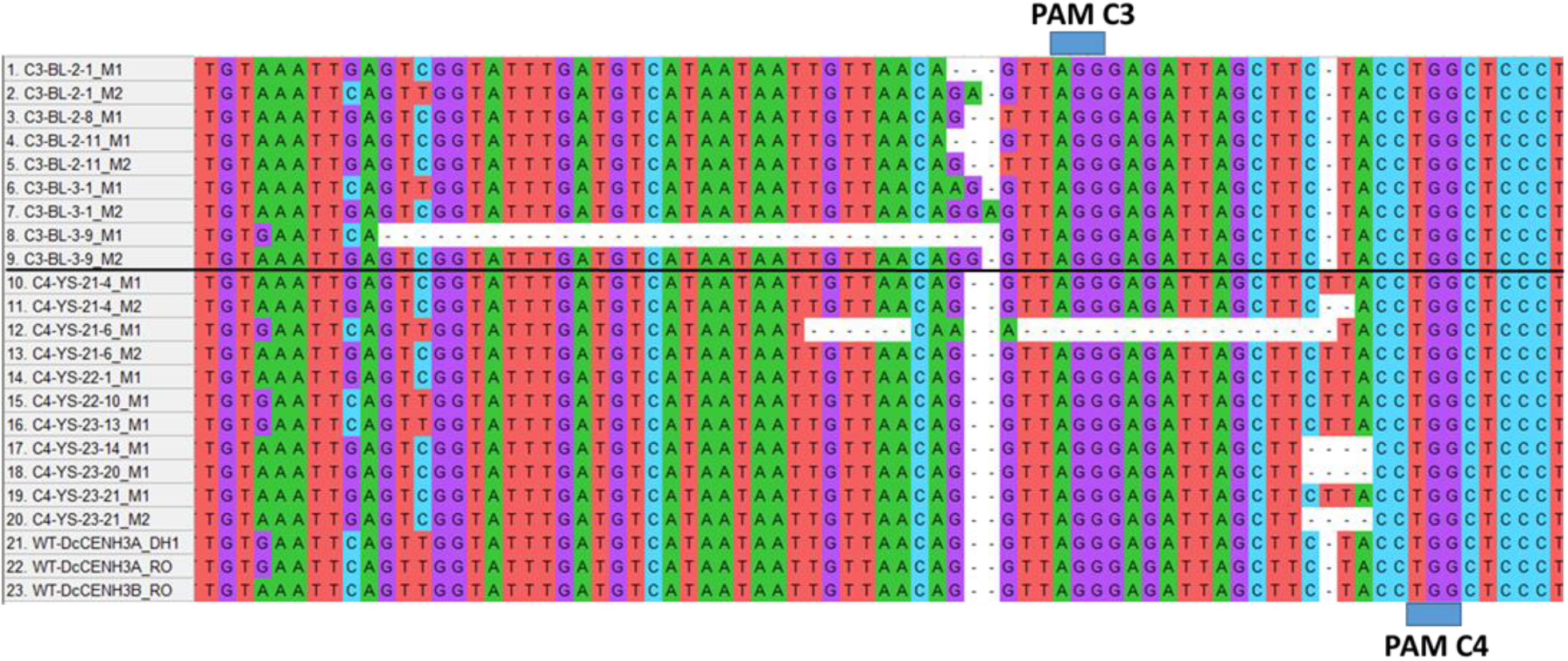
Examples for typical mutations within the genomic CENH3 sequences of regenerated T_0_ plants transformed with the C3 and C4 CRISPR constructs. Original hairy root lines were C3-BL-2 and C3-BL-3 for target C3, and C4-YS-21, C4-YS-22 and C4-YS-23 for target C4, respectively. Alignment by MUSCLE (MEGA-X).

Most mutations occurred 4 bp upstream to the PAM region. This finding was not unexpected, since the Cas9 enzyme binds to its recognition site upstream of the PAM and normally induces a DSB three bp upstream of the PAM (Schindele et al. 2018). In all plants also CENH3 wild type sequences were present at different ratios suggesting that the majority of the analysed plants were chimeric and/or heterozygous for the mutation(s). As expected, all plants showed carrot CENH3 signals after immunostaining, but in some plants the signals appeared to be somewhat weaker. Analysing the total mutation frequencies by amplicon sequencing, we noticed a tendency that the transformation with the C2 CRISPR target resulted mostly in plants with low mutation frequencies (< 10%, data not shown), when they were compared with the other two targets (C3, C4). One plant originating from the transformation of a ‘Yellowstone’ plant (C4/YS-21-6, renamed into M4-1) carrying the C4 construct displayed a 30 bp indel mutation spanning the splicing site of exon 5 (Fig. 4). Amplicon sequencing of whole transcripts obtained by RT-PCR identified a 27 bp indel mutation resulting in a translated protein sequence with the three additional amino acids F, D and V (Fig. 5). Among the about 40.000 sequences obtained after amplicon sequencing, more than 31.000 sequences showed the 27 bp indel mutation, and only less than 1.000 were wild-type CENH3 indicating that the altered transcripts might be possibly coding a functional CENH3 protein (data not shown). Up to now, all attempts to produce a flowering M4-1 mutant were unsuccessful. However, this interesting mutant could be vegetatively propagated by cuttings, which is often possible in transgenic carrot plants obtained from transformation with *R. rhizogenes*, and longer vernalization periods are currently applied to induce flowering.

**Figure 5.**
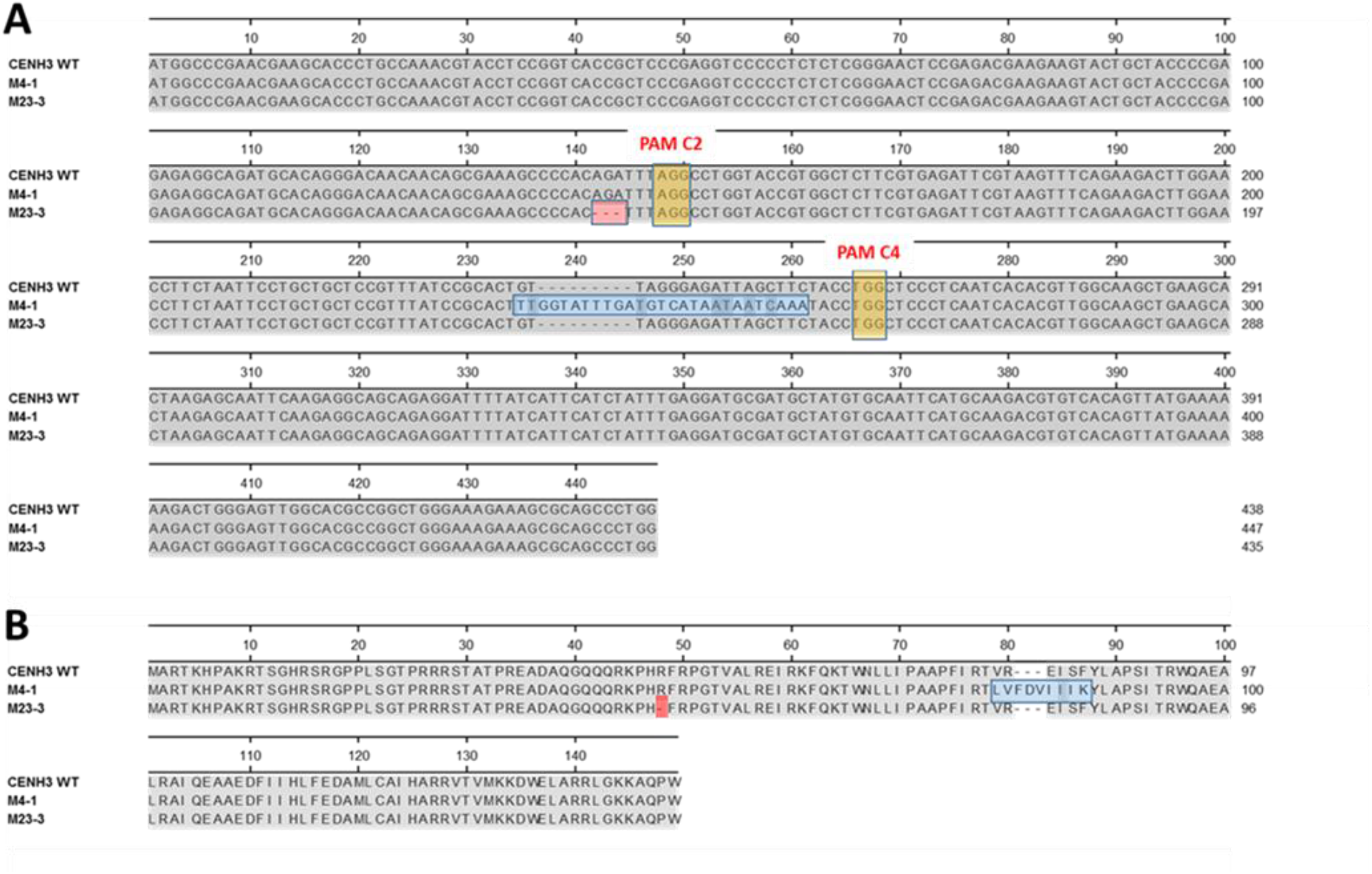
Alignment of nucleotide sequences (A) and deduced protein sequences (B) of CENH3 mutants M4-1 and M23-3

To analyse if induced mutations were heritable, and to study putative HI effects, 21 mutated T_0_ plants carrying either the C2, C3 or C4 construct were used for test crosses with carrot cultivars and progressed breeding lines, which resulted in totally 29 T_1_ progenies. Additionally, 11 T_0_ plants were used for self-fertilizations to produce 11 TI_1_ (I-inbreeding) progenies. Generally, the seed set and partly also germination was poor indicating that mutated plants were often not fully fertile. From totally 367 seeds, finally 157 plants could be raised (102 T_1_ and 55 TI_1_ plants). Flow cytometric analysis of the plants from the T_1_ generation did not indicate any haploidization effects. Amplicon sequencing was performed for the TI_1_ plants derived from selfing to prove if homozygous mutants were obtained that might be used as putative haploid inducers in a subsequent cross. Most interesting, in a single TI_1_ progeny (19.032, 10 plants derived from self-fertilization of the T_o_ plant C4/YS-21-4) a genetic segregation was found for two lethal frameshift (FS) mutations present in the T_0_ plant, either insertion ‘T’ or deletion ‘T’, respectively, and most individuals were obviously mutated on both chromosomes (Table 2). Six plants (19.032-1, −2, −4, −5, −7) with total mutation frequencies >90% showed both FS mutations with frequencies larger about 30%, and two plants (19.032-3, −6) probably were homozygous for the insertion ‘T’ mutation (>90 % insertion ‘T’). The remaining two plants (19.032-11, −14) appeared to be less frequently mutated and showed either both FS mutations (19.032-14) or only the insertion ‘T’ mutation (19.032-11). To confirm the amplicon sequencing performed by pooling we repeated sequencing for the genotypes 19.032-2, −3, −5, and −6 without usage of barcoded primer and could confirm that plants 19.032-3 and −6 were completely mutated and, therefore, were very probably homozygous for the insertion ‘T’ mutation (Table 2). Because of the lethal FS mutations, the mutated genotypes should be not able to growth. Surprisingly, the opposite was found. The plants appeared to be vital, and only flowering was delayed or suppressed. Hence, contrarily to the assumption of a one-gene control, this finding suggested the existence of a second CENH3 gene in the carrot genome.

### Identification of a second CENH3 gene in the carrot genome

Based on bioinformatic searches in the carrot genome vers.2 (Iorizzo et al. 2016) we found an additional sequence (DCAR_021490) on chromosome 6, which appeared to encode a second CENH3 gene. This putative CENH3 gene was only fragmentarily assembled and annotated and, therefore, has not been considered yet in the carrot CENH3 research project. The CENH3 gene, which has been initially the target gene for the CRISPR/Cas9 approach of this study, is located on carrot chromosome 7. This gene previously isolated and functionally characterized by Dunemann et al. (2014) could be in principal confirmed by the carrot genome sequence, but the annotation of this gene (DCAR_025246) was not complete. The genomic sequence of DCAR_025246 is about 800 bp shorter as the whole genomic sequences of the CENH3 gene previously cloned from ‘Deep Purple’ (DP) and a homologous CENH3 gene cloned from ‘DH1’ in this study. In addition, the CDS and the translated protein sequence of DCAR_025246 contained two gaps at the beginning (amino acids 19-35) and at the end (Fig. 6). We re-analysed this genomic region and noticed that this CENH3 gene was indeed completely present in the carrot genome. An optimized prediction was performed and called DCAR_025246-OP. The genomic position of the DNA sequence of DCAR_025246-OP in the actual carrot genome is on chromosome 7 from 25.166.681 to 25.171.180 Mbp. With regard to the second putative CENH3 gene on chromosome 6, the corresponding DCAR_021940 sequence has a total length of about 11 kb, but only the first 1.000 bp could be partly assigned to CENH3. As shown in Figure. 6, less than half of the coding sequence is present in DCAR_021940. Attempts to find the missing parts in this genomic region failed. Interestingly, the same gap (PPL….TPR) is present in both DCAR_021940 and DCAR_025246, probably due to the same annotation problem.

**Figure 6.**
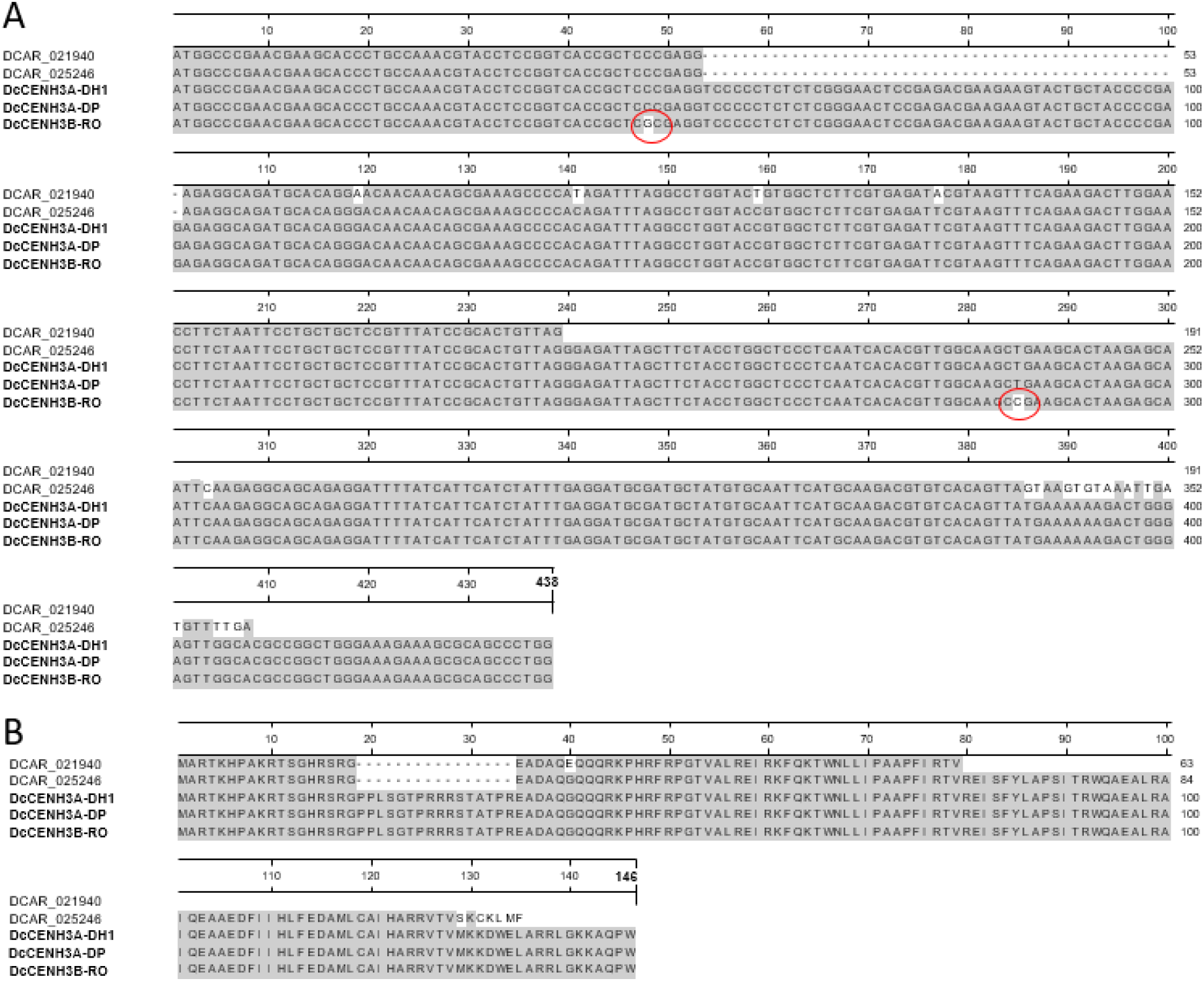
Alignment of CDS (**A**) and translated protein sequences (**B**) of annotated CENH3 gene models DCAR_021940 and DCAR_025246 present in the carrot genome vers.2 (Iorizzo et al. 2016) and three carrot CENH3 genes cloned from DH1, DP and RO. SNPs are marked with a red circle.

Based on sequence differences between the two DCAR sequences within the first 1.000 bp of the genomic sequences, primer pairs were designed to prolong step-by-step the unknown (about 3.500 bp expected) part of the DCAR_021940 gene by PCR. Forward primer specific for DCAR_021940 were combined with unspecific primer developed from the available DCAR_025246 sequence. If amplicons of the expected size were obtained, they were cloned and sequenced, and the next primer pair was designed. By using this cloning strategy, we were finally able to develop a PCR primer pair from the 5’UTR and 3’UTR regions for cloning the complete full-length genomic DNA sequence from cultivar ‘Rotin’ (RO). This new gene on carrot chromosome 6 was designated as *DcCENH3B*, and the gene corresponding to DCAR_025246 on chromosome 7 was given the name *DcCENH3A*. By using another specific forward PCR primer based on the first nt’s of the CDS of DCAR_025246, we cloned the genomic sequence of *DcCENH3A* from the ‘DH1’ genotype, which was used for carrot genome sequencing by Iorizzo et al. (2016). The genomic sequence of *DcCENH3B* of ‘Rotin’ (*DcCENH3B-RO*) has a length of 4.430 bp, and those of *DcCENH3A* from ‘DH1’ (*DcCENH3A-DH1*) is 4.500 bp long, respectively (Suppl. Fig. 1; Suppl. Inf. 1). An alignment of the two newly cloned genes, the earlier cloned CENH3 gene from ‘Deep Purple’ (with the name *DcCENH3A-DP*), and DCAR_025246-OP is shown in Suppl. Fig. 1. The sequence of the cloned gene *DcCENH3A-DH1* is to 100% identical with the optimized prediction of DCAR_025246-OP (Suppl. Fig. 2). *DcCENH3B-RO* displayed 94 to 95% sequence identity with the two CENH3A genes. The coding sequences, which are only about one tenth as long as the genomic sequences, however, are very similar. Only two SNPs were found in the CDS of *DcCENH3B-RO*, which does not change the protein sequence (Fig. 6). The nucleotide sequence lengths of 438 bp, encoding a 146 amino acid protein, are identical for all three cloned CENH3 genes. Based on the carrot genome sequence vers.2 implemented in *Phytozome v.13* (Goodstein et al. 2012) and *PlantCARE*, a database of plant cis-acting regulatory elements (Rombauts et al. 1999), we bioinformatically identified the putative promotor sequences of *DcCENH3A* on chromosome 7 and *DcCENH3B* on chromosome 6 (Suppl. Inf. 2). An analysis of motifs encoding putative regulatory elements is shown in Suppl. Fig. 3.

The finding that carrot obviously possess duplicated CENH3 genes on chromosomes 6 and 7 raises the question about the functional roles of both CENH3 paralogs. In most diploid eukaryotes and flowering plant species, CENH3 is encoded by a single copy gene (Ishii et al. 2020). Knowledge about duplicated CENH3 genes is restricted to a small number of about a dozen plant species. Two CENH3 genes were reported for *Arabidopsis halleri* and *A. lyrata* (Kawabe et al. 2006), *Hordeum vulgare* (Sanei et al. 2011), *Pisum sativum* (Neumann et al. 2012), *Luzula nivea* (Moraes et al. 2011), *Lathyrus sativus* (Neumann et al. 2015), *Secale cereale* (Evtushenko et al. 2021) and *Vigna unguiculata* (Ishii et al. 2020). In genomes carrying a gene duplication, often one of them is silenced, but sometimes both duplicated genes are expressed (Moraes et al. 2011). In tetraploid *Oryza* species, two CENH3 genes were identified, and both are transcribed, showing no preferential expression of one of them (Hirsch et al. 2009). With regard to the putative functions of the duplicated CENH3 genes in carrot, we could not yet verify if both genes are functional. Because of the 100% identical protein sequences, it is impossible to create a specific polyclonal antibody for functional analysis of *DcCENH3B*. In addition, gene-specific RT-PCR was not possible yet. An alignment of the deduced proteins of the duplicated CENH3 genes found in the literature indicated that the two carrot CENH3 genes are most similar, which suggests an early evolutionary origin (alignment not shown). We used a universal PCR primer pair from the beginning and the end of the CDSs of *DcCENH3A* and *DcCENH3B* to amplify a 437 bp transcript, which was analysed by Illumina amplicon sequencing. In addition, amplicons were cloned and analysed by plasmid sequencing. Up to now, we found no indications that the investigated plants from cultivars ‘Rotin’, ‘Yellowstone’ and ‘DH1’ expressed the *DcCENH3B* gene. All transcripts were *DcCENH3A*, using the detected two SNPs as criterion. Interestingly, in the “lethal” CENH3 mutants of the TI_0_ progeny 19.032 described before, only wild-type *DcCENH3A* appeared to be expressed, and analyses of the genomic sequences flanking the C4 CRISPR target site indicated that *DcCENH3B* was that gene with the induced knockout mutations. In contrast, the M4-1 mutant showing almost exclusively mutated (27 bp indel) transcripts appeared to be mutated within the *DcCENH3A* gene. However, it must be stated that a safe verification of the targeted gene only by using the short genomic sequences (370 to 400 bp) obtained by amplicon sequencing was not possible in this study. Carrot is a highly heterozygous species, and it appeared that there is considerable allelic variability for CENH3 not only between cultivars. Even on the two homologous chromosomes of a distinct carrot genotype allelic variability was observed. This complex genetic situation, together with the challenge to sequence different target regions for two CENH3 genes, makes a solid sequencing study for characterization of hundreds of putative mutated plants to a practically unsolvable task. In addition, each sequencing study is only a snapshot in time for a specific genotype. As far the CRISPR targets are still present in the plant genome, as it is often the case after crossing, new mutations can occur in the next generation, which would only be detected after repeated sequencing. Since the three CRISPR targets C2, C3, and C4 have been cloned before the detection of *DcCENH3B*, we wanted to know, if the constructs would be able to target also this gene. As shown in Suppl. Fig. 4, this seems to be case. Nevertheless, the knowledge about the existence of a duplicated CENH3 locus in carrot implicates that the creation of a HI plant based on the 1-STEP approach might be difficult, if not impossible. This option would probably only function, if *DcCENH3B* is permanently inactive. More basic research would be needed on the function of both CENH3 genes in carrots, to justify further 1-STEP attempts.

### 2-STEP experiments

First experiments to use *R. rhizogenes* for co-transformation of the CRISPR C4 target construct and the ginseng *PgCENH3* gene construct have been started parallel to the 1-STEP experiments (without prior knowledge about the second CENH3 gene on chromosome 6), and preliminary results were reported (Dunemann et al. 2019). With this approach, we wanted to use co-transformation to develop an accelerated 2-STEP HI system by combining mutation induction with the simultaneous integration of the foreign CENH3 gene provided for complementation of lethal mutations in the endogenous gene. The used “rescue” gene was *PgCENH3* cloned by a PCR approach from transcripts of *Panax ginseng*, a member of the plant family *Araliaceae* (order Apiales). *PgCENH3* has exactly the same length as the CENH3s of *D. carota* (146 amino acids) but displays more than 30 amino acid changes especially within the N-terminal region of the gene (Fig. 3). Initially a test transformation only with the *PgCENH3* vector was performed, and it was shown, that *PgCENH3* was stably expressed in transgenic carrots and the ginseng CENH3 protein co-localized with carrot CENH3 (Fig. 7). Co-transformation together with the binary vector carrying the C4 CRISPR target construct resulted in totally 50 regenerated T_0_ plants, which were analysed by PCR using the SS42/43 and *bar* primer (C4 target) and the PCEN primer (ginseng CENH3) to verify the double-transformation with both constructs. In comparison to the 1-STEP experiment targeting C4, the same spectrum of mutations was found in the C4 target region, but it seemed that the frequency of strong mutations (frameshift mutations) was lesser, and the frequency of weak mutations (silent mutations due to single nt substitutions) and wild-type plants was higher than in the 1-STEP experiment. In addition, plants/tissues appeared also here to be often strongly chimeric concerning the CRISPR effects. As in the 1-STEP approach, some selected T_0_ mutants carrying the ginseng CENH3 gene were used to generate the TI_1_ generation by self-pollination (to get homozygous mutants) and T_1_ progenies by crosses with carrot cultivars (to study putative HI effects). As expected, *PgCENH3* segregated in the next generation, but the genetic transmission of the C4 target mutations appeared to be generally low. To screen efficiently several hundreds of progeny plants for putative reduction or failure of carrot CENH3 signals, we used a slightly modified immunostaining protocol, which allowed a fast evaluation due to a clustering of stained nuclei (Fig. 7). Only in a few cases, as for example in the T_0_ individual G4/YS-1-4, mutations were possibly associated with a reduced carrot CENH3 protein accumulation, but no plant was found with a complete failure of carrot CENH3, neither in the T_o_ nor in the T_1_ and TI_1_ generations. Within the T_1_ progeny 18.021 deriving from a cross with G4/YS-1-4 as mother, flow cytometry, chromosome counting based on root tip mitosis, and CENH3 immuno-fluorescence staining showed irregularities in cell divisions in leaves of young seedlings. One plant (18.021-1) appeared to be partially haploid (Suppl. Fig. 5). In addition, molecular analyses showed that all transgenes were lost in the plants of this progeny, which might be a sign that the genome of the putative HI might have been eliminated. However, a repeated cytogenetic analysis based on older plants could not confirm the existence of any homogenously haploid plant.

**Figure 7.**
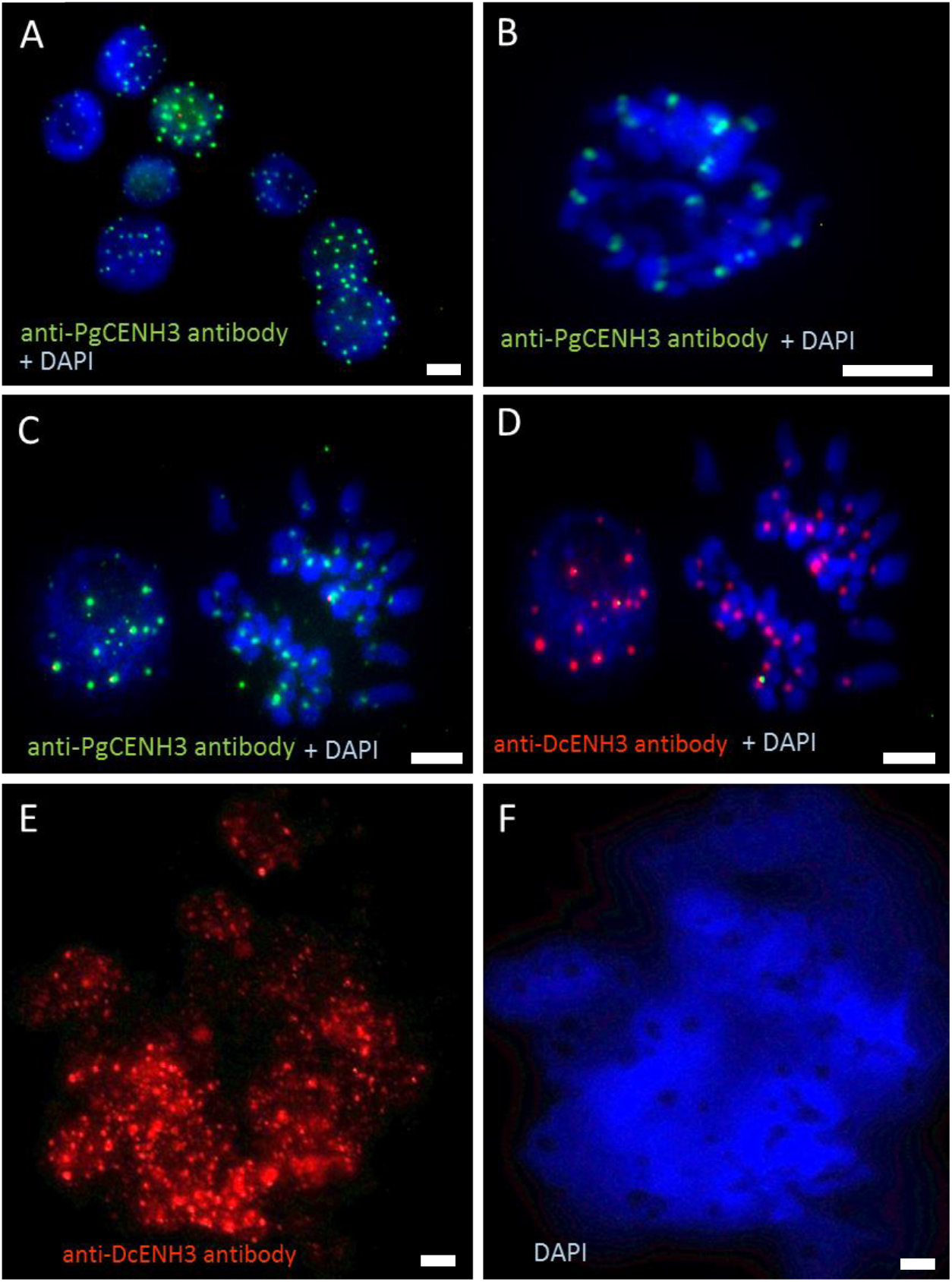
(**A-B**) Immunostaining of a nuclei preparation from young leaves of a carrot plant trans-formed with the *PgCENH3* gene using anti-*PgCENH3* antibody (A) and immunostaining of metaphase chromosomes of a root-tip cell of the same plant using anti-*PgCENH3* antibody (B). (**C-D**) Double-immunostaining of a root tip preparation of a ginseng-CENH3-transformed carrot plant with both anti-*PgCENH3* antibody (C) and anti-*DcCENH3* antibody (D).(**E-F**) Immunostaining of nuclei clusters prepared from young leaves of a putative CENH3 mutant with anti-*DcCENH3* antibody (E) and DAPI staining of the same preparation (F). Scale bar 5 μm.

Because of the obviously low mutation efficiency and the poor transmission of mutations in the next generations we set up a new crossing experiment with the aim to combine ginseng CENH3 producers with the strongest carrot CENH3 mutants identified in the 1-STEP approach. Seven 2-STEP T_1_ plants with strong expression of ginseng CENH3 (and carrot CENH3 wild-type after immunostaining), which still possessed the C4 CRISPR target construct, were selected for crosses with T_0_ plants out of the 1-STEP approach. These plants have been selected because of their amplicon sequencing results, which have revealed putative lethal (heterozygous) mutations induced either by the C3 or C4 constructs. Surprisingly, in the sequenced 33 progeny plants the mutation frequencies decreased, and only in the progenies 19.300-4 and 19.300-8 the frequencies were in the magnitude of the 1-STEP parent plant (Table 3). The reason for the obvious decrease of the mutation frequencies is unknown. One reason might be, that strong CENH3 mutations did not pass the meiose due to occurrence of lethal alleles. Why could the ginseng CENH3 stably present in a T_1_ plant obviously not complement the lethal carrot CENH3 mutations? Since the ginseng CENH3 is controlled by a doubled p35S promotor, it cannot be ruled out that the activity of *PgCENH3* was absent or too low during the gametogenesis phase. In *A. thaliana*, CENH3-GFP expression under the control of p35S promotor resulted in centromere specific GFP signals in a large number of organ types such as root tips, leaf meristems, young leaf material, flower meristems and developing flower organs, but no expression was observed in male and female reproductive cells or in developing meiocytes (De Storme et al. 2016). However, in strawberries p35S appeared to drive expression of GUS in pollen and other reproductive tissues (De Mesa et al. 2004). Although it is a generally held view that the 35S promotor is virtually silent in reproductive tissues, Dutt et al. (2014) concluded that the transgene expression profile conferred by the CaMV 35S promotor is species-dependent and may show variability due to its interaction with environmental factors and the physiological state of the plant. One option for future carrot 2-STEP approaches might be the usage of a construct combining the native *DcCENH3A* promotor with the CDS of *PgCENH3*.

**Table 3.** Mutation frequencies evaluated by amplicon sequencing of the C3 and C4 target regions of parental individuals and progeny plants derived from crosses between 1-STEP and 2-STEP plants

| Progeny Name | Mother | Target and Origin | Mutated sequences (%) | Father | Target and Origin | Mutated sequences (%) | No. of sequenced progeny plants | Mutated sequences (%) Range |
| --- | --- | --- | --- | --- | --- | --- | --- | --- |
| 19.200-1 | C4/YS-22-8 | C4 (1-STEP) | 60 | 18.005-10 | C4 (2-STEP) | 7,2 | 1 | 10 |
| 19.200-2 | 18.005-10 | C4 (2-STEP) | 7,2 | C4/YS-22-8 | C4 (1-STEP) | 60 | 4 | 5,7 - 18,2 |
| 19.200-4 | C4/YS-23-9 | C4 (1-STEP) | 21,6 | 18.006-17 | C4 (2-STEP) | 7,8 | 1 | 6,5 - 6,7 |
| 19.200-5 | C4/YS-23-11 | C4 (1-STEP) | 19,7 | 18.006-17 | C4 (2-STEP) | 7,8 | 5 | 5,4 - 6,5 |
| 19.200-7 | 18.003-10 | C4 (2-STEP) | 15,7 | C4/YS-23-11 | C4 (1-STEP) | 19,7 | 3 | 6,5 - 12,8 |
| 19.200-8 | 18.005-10 | C4 (2-STEP) | 7,2 | C4/YS-23-11 | C4 (1-STEP) | 19,7 | 1 | 4,8 - 5,4 |
| 19.200-9 | C4/YS-23-12 | C4 (1-STEP) | 7 | 18.006-17 | C4 (2-STEP) | 7,8 | 3 | 6,0 - 8,1 |
| 19.300-3 | C3/BL-5-3 | C3 (1-STEP) | 16,2 | 18.006-17 | C4 (2-STEP) | 7,8 | 9 | 6,4 - 13,1 |
| 19.300-4 | 18.001-32 | C4 (2-STEP) | 6,5 | C3/BL 5-5 | C3 (1-STEP) | 20,6 | 4 | 17,9 - 21,9 |
| 19.300-8 | C3/BL-3-8 | C3 (1-STEP) | 34,1 | 18.005-14 | C4 (2-STEP) | 8,3 | 1 | 26,2 |
| 19.300-9 | C3/BL-3-9 | C3 (1-STEP) | 34 | 18.005-10 | C4 (2-STEP) | 7,2 | 1 | 11,9 |

Because of the low mutation frequencies found in generative descendents we wanted to know, if combinations of different CRISPR/Cas9 targets would enhance the chance to generate more strong mutations able for HI induction already in the T_0_ generation. For this approach, we used the same carrot T_1_ plant material used for the crosses with 1-STEP mutants and used individuals with proven strong presence of ginseng CENH3 for a second transformation. In this 2-STEP-DT experiment combinations of the two CRISPR targets C2 and C3 or C2 and C4, respectively, were introduced by co-transformation in the transgenic carrots carrying the ginseng CENH3 gene and the C4 target too. The mutation spectrum found after amplicon sequencing for each target region in the transformants was comparable with earlier experiments, with the difference, that more interesting mutations were found in the C2 target than in the 1-STEP experiment (Fig. 8). PCR with a universal primer pair (DEL-C2C4-F/-R) flanking the C2 and C3/C4 targets of both CENH3 genes, however, did not give any incidence, that large deletions (i.e. between the C2 and C4 targets) could be induced. Among about 60 sequenced T_0_ plants, 15 plants were found which contained strong mutations (different types of insertions and deletions) at different frequencies. Both *DcCENH3A* and *DcCENH3B* appeared to be concerned, but as mentioned before, a detailed quantification was not possible due to additional SNPs present in the target regions of this genetically diverse material. In some mutants, mutation frequencies of up to 40% indicated heterozygous mutations. The most interesting finding was a putative homozygous in-frame mutation in *DcCENH3A* of mutant M23-3 concerning a 3-bp deletion in target C2, which results in the deletion of a single arginine (’R’) amino acid just before before the αN-helix (Fig. 5, Suppl. Fig 6). Among more than 90.000 sequences obtained after transcript amplicon sequencing, no wild-type sequences were found indicating that the altered CENH3 transcripts might be functional. Interestingly, in the study of Kuppu et al. (2020) CRISPR/Cas9-mediated in-frame deletions at the beginning of the highly conserved αN-helix of the CENH3-HFD such as a 2-amino acid (-GT) mutation shown in Suppl. Fig. 6 were able to efficiently induce haploids in *A. thaliana*. Unfortunately, original mutant M23-3 as well as vegetatively propagated cuttings have not yet flowered. Plant growth is very slow suggesting that the “minus R” mutation might have some serious effects on cell division.

**Figure 8.**
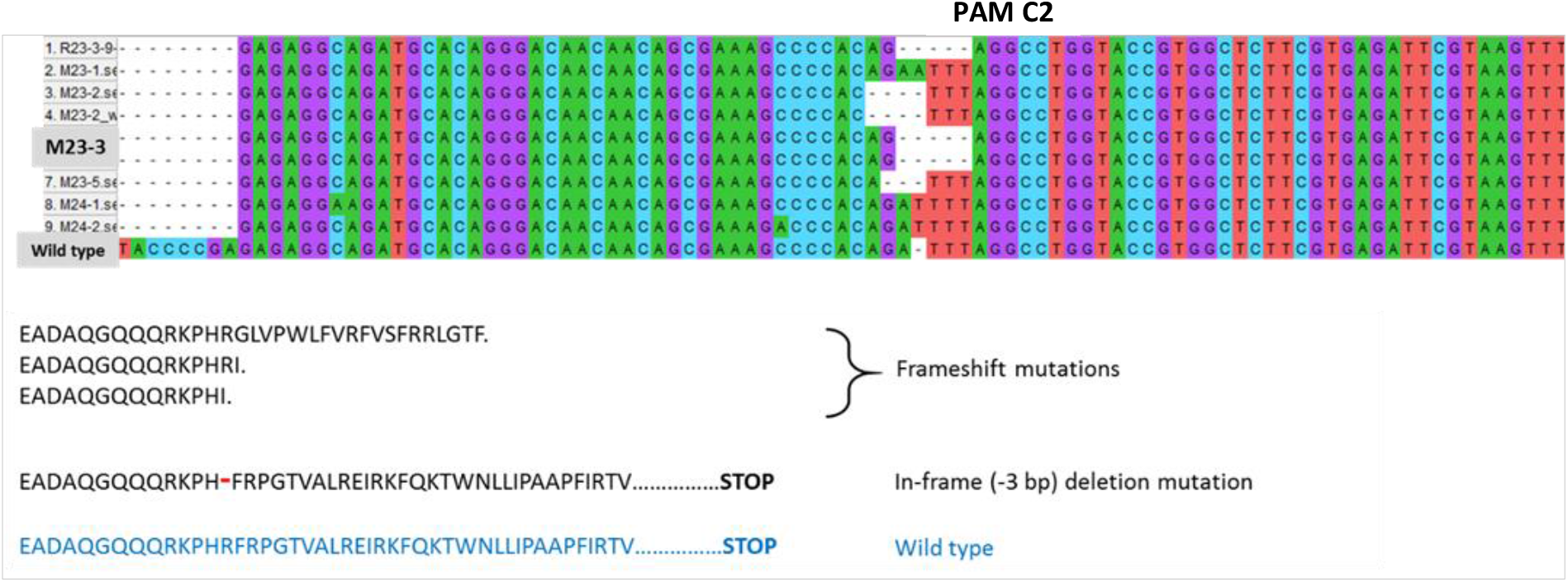
Three different frameshift mutations and one in-frame mutation in C2 target sequences of T_0_ plants derived from the double-transformation experiment

## Conclusion

Several transformation experiments and subsequent genetic experiments have shown that CRISPR/Cas9-based gene editing of carrot CENH3 is feasible. Promising mutants such as the 27-bp in-frame indel mutant (1-STEP approach) or the 3bp in-frame deletion mutant (2-STEP approach) have been found, but their successful usage as putative haploid inducers is uncertain yet. The finding of a duplicated CENH3 locus in the carrot genome would require more research on the functional roles of both genes to develop the best strategy for future CENH3 gene editing.

Next generation sequencing of amplicons spanning CRISPR target sites and transcript-based amplicon sequencing seemed to be appropriate methods to select promising mutants, to estimate mutation frequencies, and to allow a first prediction which CENH3 gene was concerned. Since there was a serious amount of allelic variability within CENH3 genes of different carrot cultivars, future carrot CENH3 research should consider cloning and sequencing of further full-length CENH3 genes. A more practical alternative might be to use homozygous carrots, such as double-haploid carrot lines, for targeting CENH3 and to analyse mutants by deep sequencing.

The main lesson from the 2-STEP experiments is that a simultaneous carrot CENH3 gene editing by CRISPR/Cas9 and integration of a foreign CENH3 gene, in this case from *Panax ginseng*, seemed to be easily possible by co-transformations based on *R. rhizogenes*. The ginseng CENH3 protein was stably accumulated inside the kinetochore region of carrot chromosomes, indicating that *PgCENH3* might be a suited candidate for complementation of putative carrot CENH3 knockout mutants. However, presently it is still unclear, if this gene, controlled by a CaMV 35S promotor, is fully functioning during the meiotic cell divisions and, therefore, suited to complement lethal gametes. Usage of a native carrot CENH3 promotor might improve CRISPR mutation accumulation in future 2-STEP approaches.

## Acknowledgements

We wish to thank Thorben Sprink (JKI Quedlinburg) for cloning the carrot CENH3 targets into the pDE-Cas9 vector and Götz Hensel (Heinrich-Heine-University Düsseldorf) for advices in cloning the ginseng CENH3 construct. We also thank Katrin Kumke and Andreas Houben (IPK Gatersleben) for help in carrot immunofluorescence assays.

## Supplemental Tables 1 and 2

**Suppl. Table 1.** Used media for *R. rhizogenes* cultivation and transformation

| Medium | Basic components | Mineral nutrients | Antibiotics / Selective agents | Usage |
| --- | --- | --- | --- | --- |
| CPY-S1 | bacto-yeast extract 0,1% (Difco),<br>peptone 0,5% (Roth),<br>saccharose 0,5% (Merck),<br>bacto-agar 20g/l (BD) | MgSO<sub>4</sub>·7H<sub>2</sub>O<br>200 mM | spectinomycin 100 mg/l<br>(Sigma-Aldrich) | culture of *R. rhizogenes*-strains |
| CPY-S2 | bacto-yeast extract 0,1% (Difco),<br>peptone 0,5% (Roth),<br>saccharose 0,5% (Merck) | MgSO<sub>4</sub>·7H<sub>2</sub>O<br>200 mM | spectinomycin 10 mg/l<br>(Sigma-Aldrich) | culture of *R. rhizogenes*-strains |
| WA-CC | agar 2 g/l (Duchefa) |  | cefotaxime 200mg/l (Duchefa),<br>carbenicillin 100 mg/l (Roth) | transformation / regeneration |
| MS-CCP | plant agar 6 g/l (Duchefa),<br>saccharose 20 g/l (Merck) | Murashige & Skoog medium<br>(micro and macro elements incl. vitamins<br>(Duchefa M0222)) | cefotaxime 200mg/l (Duchefa),<br>carbenicillin 100 mg/l (Roth),<br>phosphinotricin 10 mg/l (glufosinate-ammonium pestanal)<br>(Sigma-Aldrich) | hairy root growth / selection |

**Suppl. Table 2.**
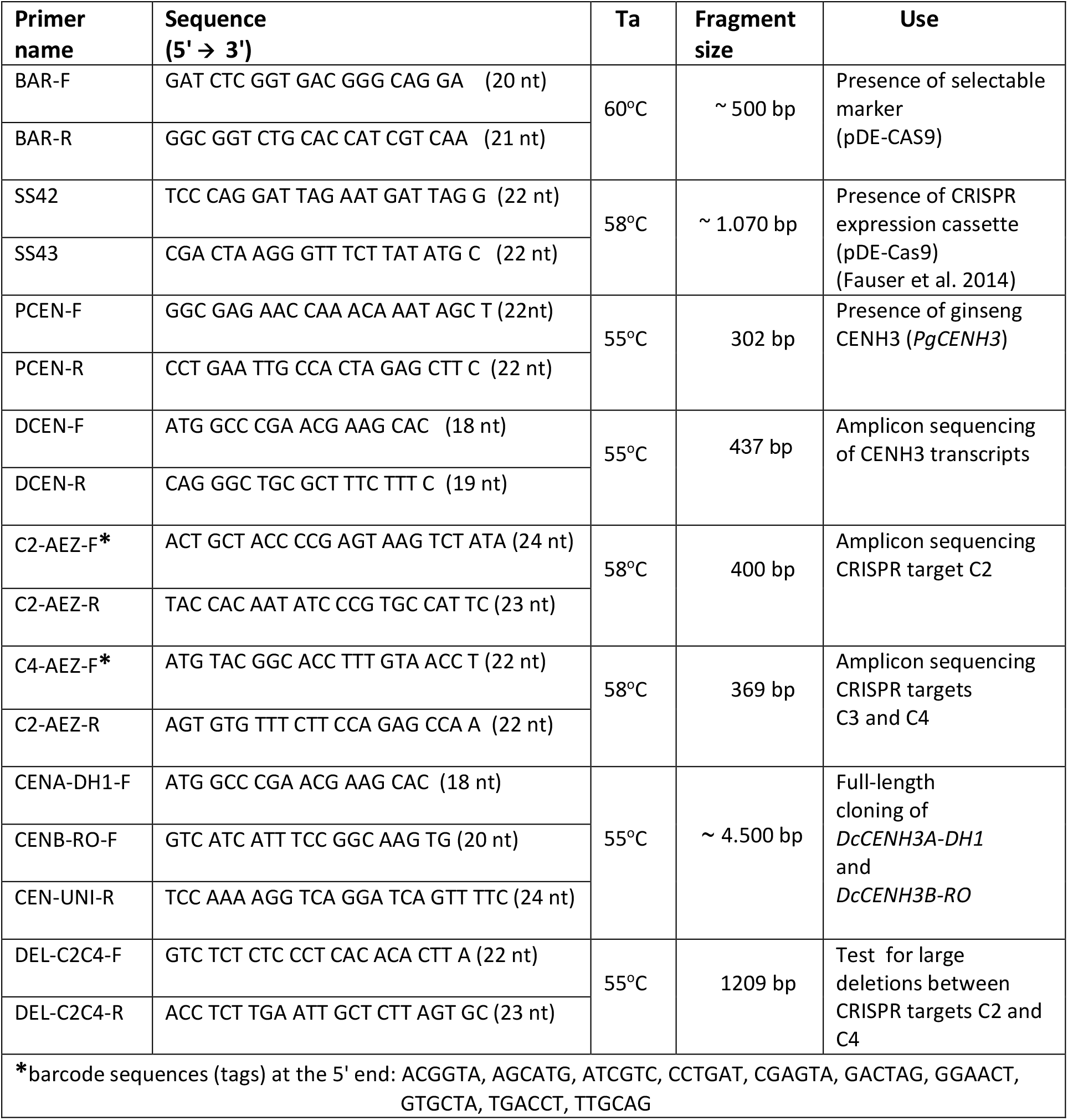
PCR primer information

## Supplemental Figures 1 - 6

**Suppl. Figure 1.**
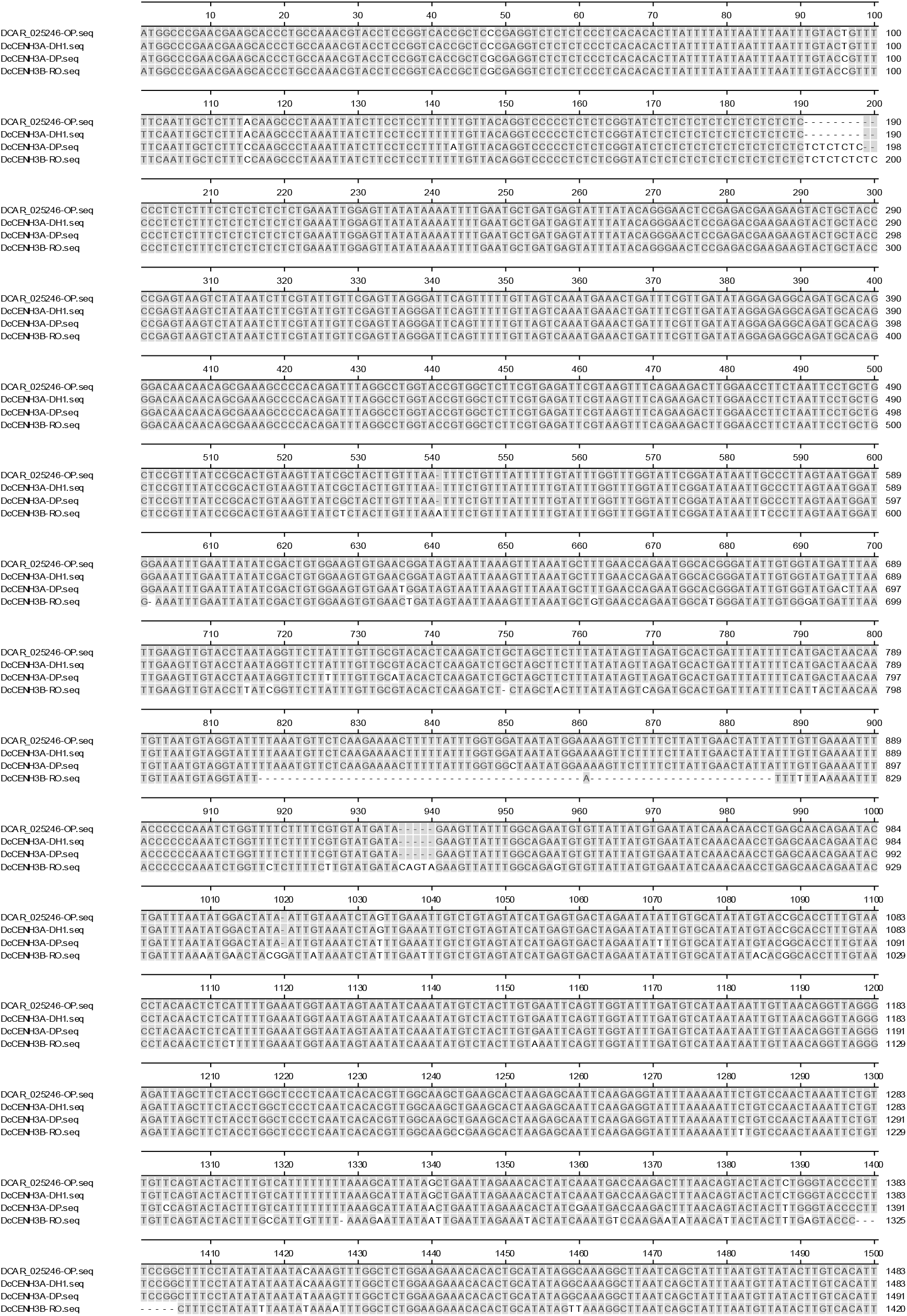
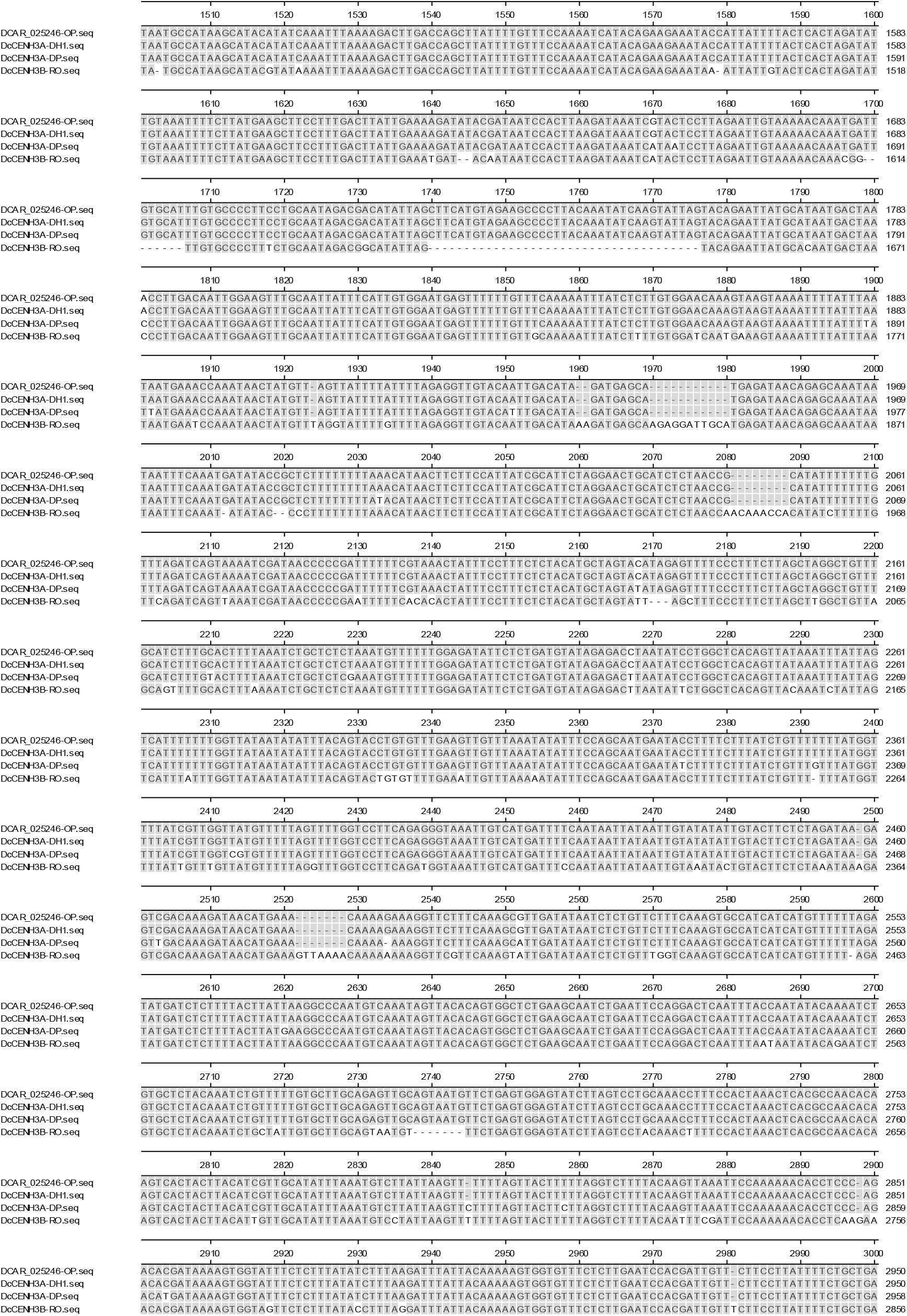
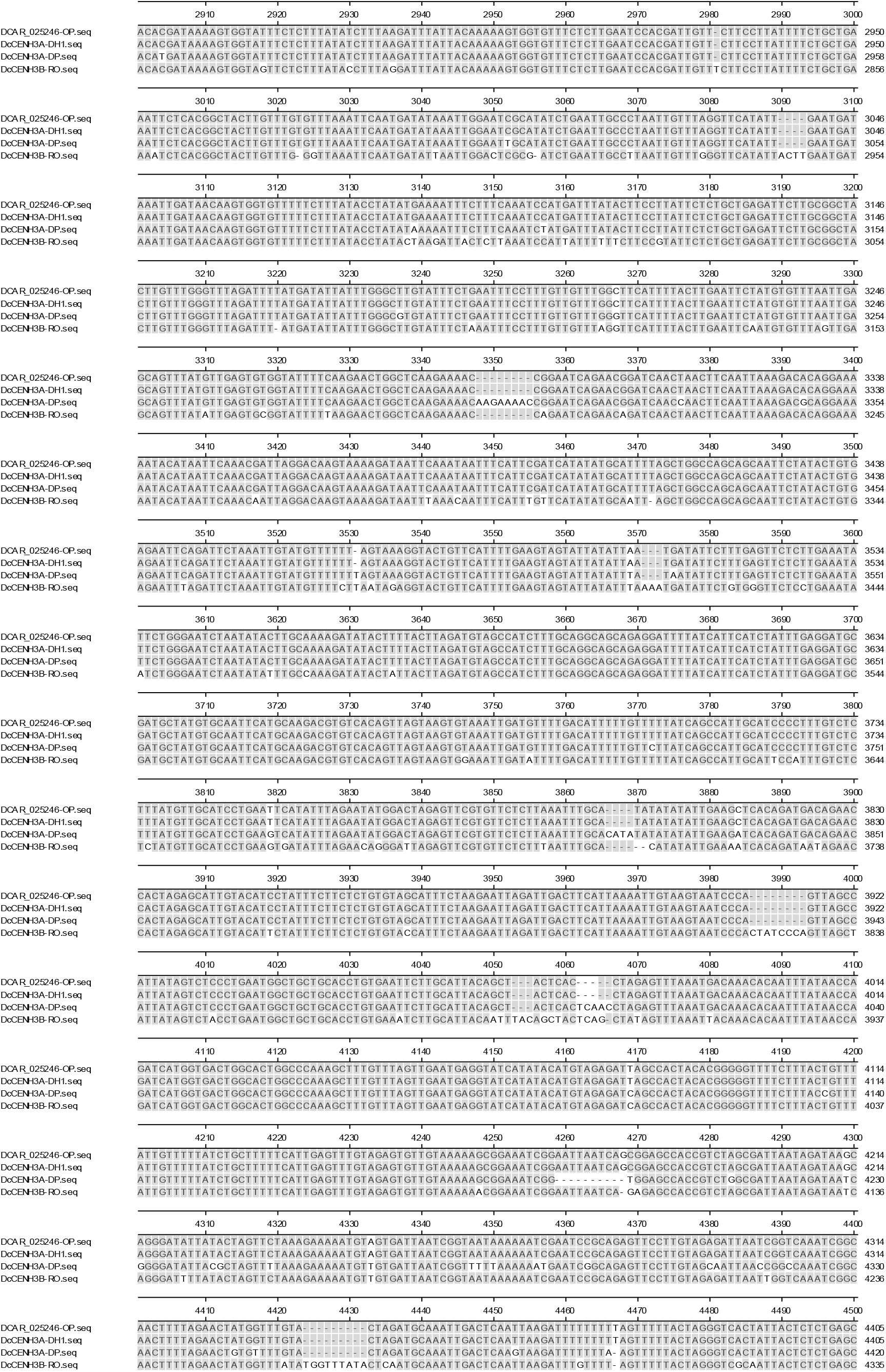
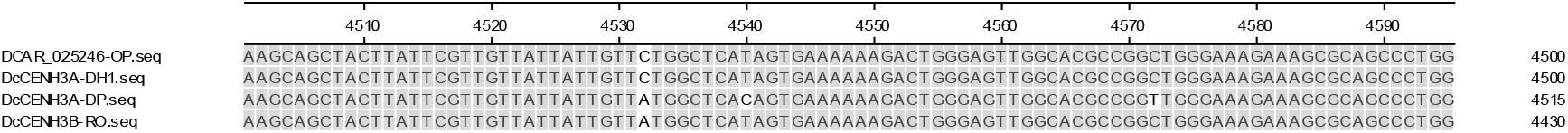
Alignment of full-length genomic DNA sequences of two carrot CENH3A genes cloned from DP and DH1, and *DcCENH3B-RO* (DNASTAR Lasergene, ClustalW)

**Suppl. Figure 2.**
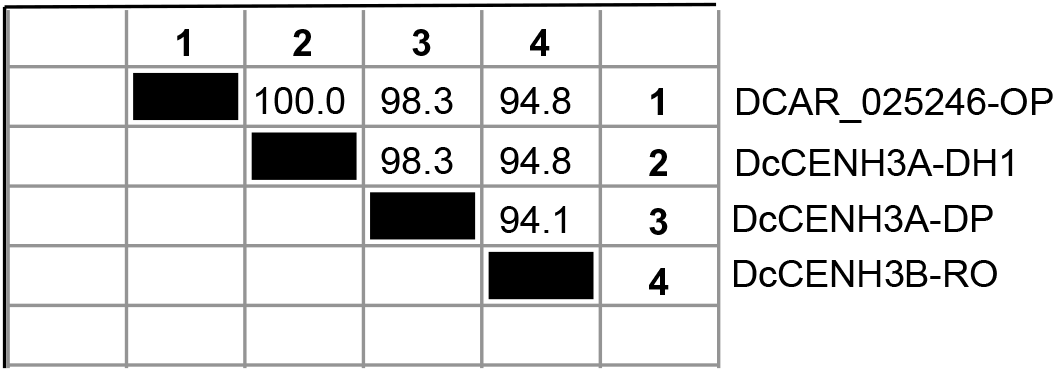
Nucleotide sequence distances (% identity) of genomic sequences of *CENH3* genes cloned from DH1, DP and RO and the optimized prediction of DCAR_025246. For alignment, see Suppl. Fig. 1

**Suppl. Figure 3.**
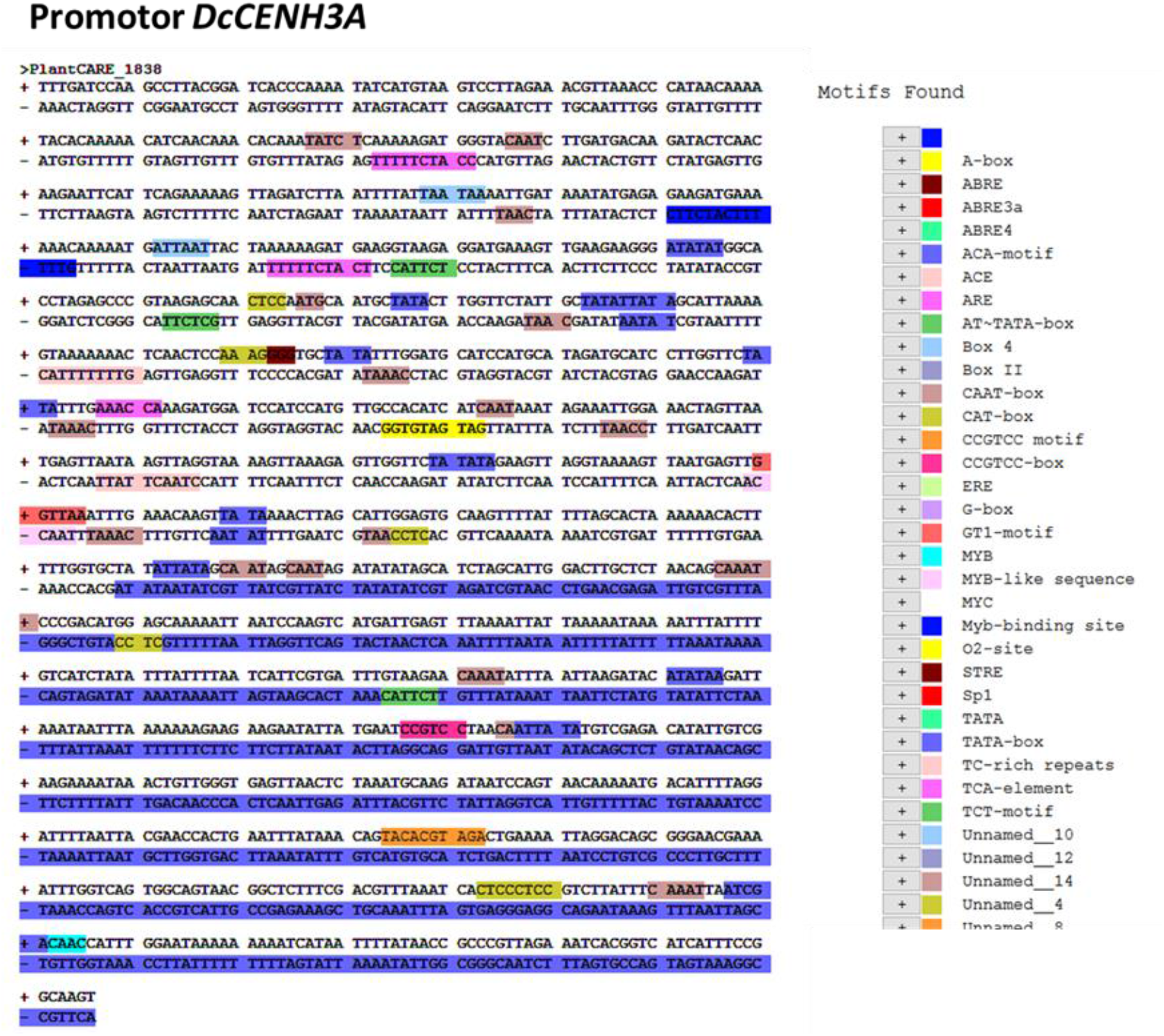
(A) Analysis of putative *DcCENH3A* promotor using *PlantCare* database.

**Suppl. Figure 3.**
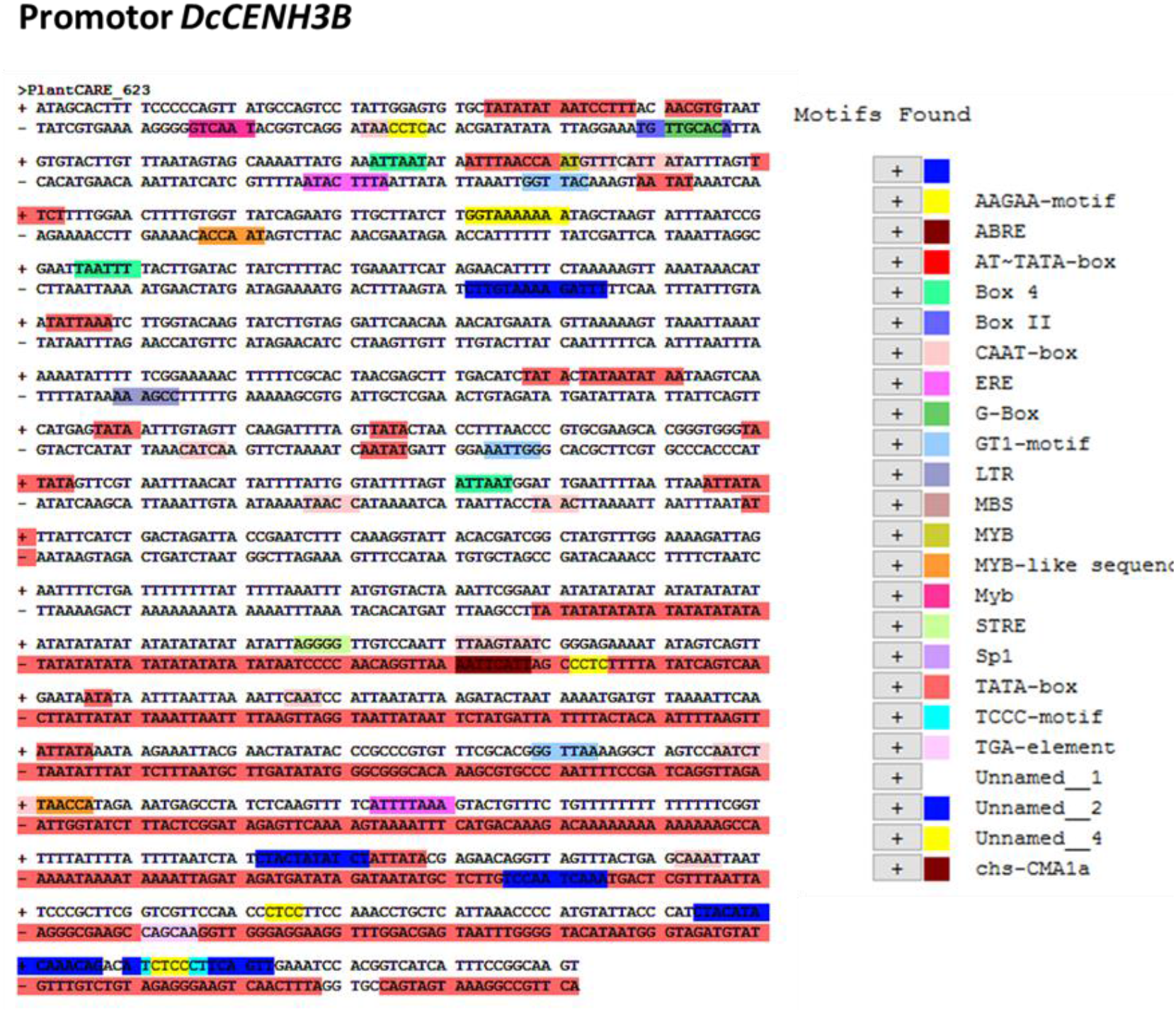
(B) Analysis of putative *DcCENH3B* promotor using *PlantCare* database

**Suppl. Figure 4.**
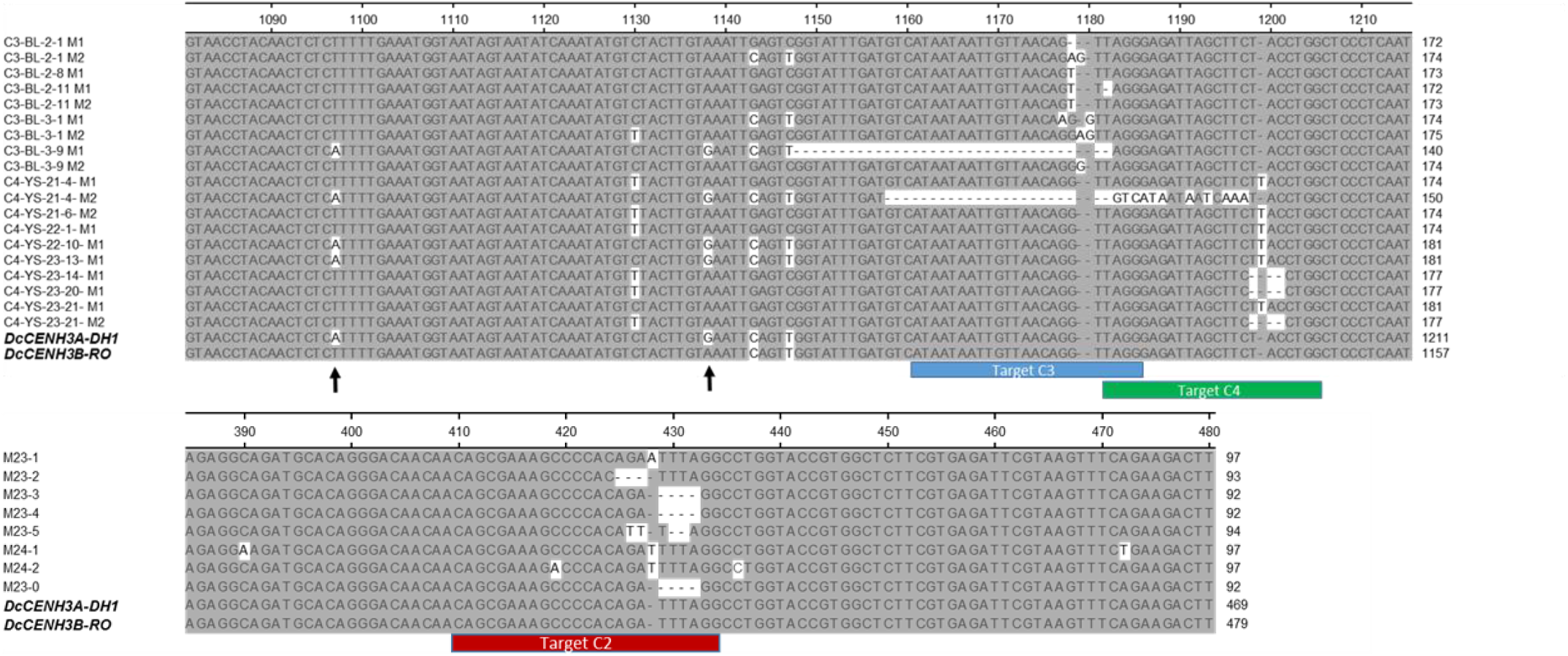
Location of the CRISPR targets C3 and C4 (top) and target C2 (bottom) in the genomic sequences of newly cloned CENH3 genes compared with sequences obtained by amplicon sequencing of some mutants. Arrows show SNPs present in the CENH3A and CENH3B genes.

**Suppl. Figure 5.**
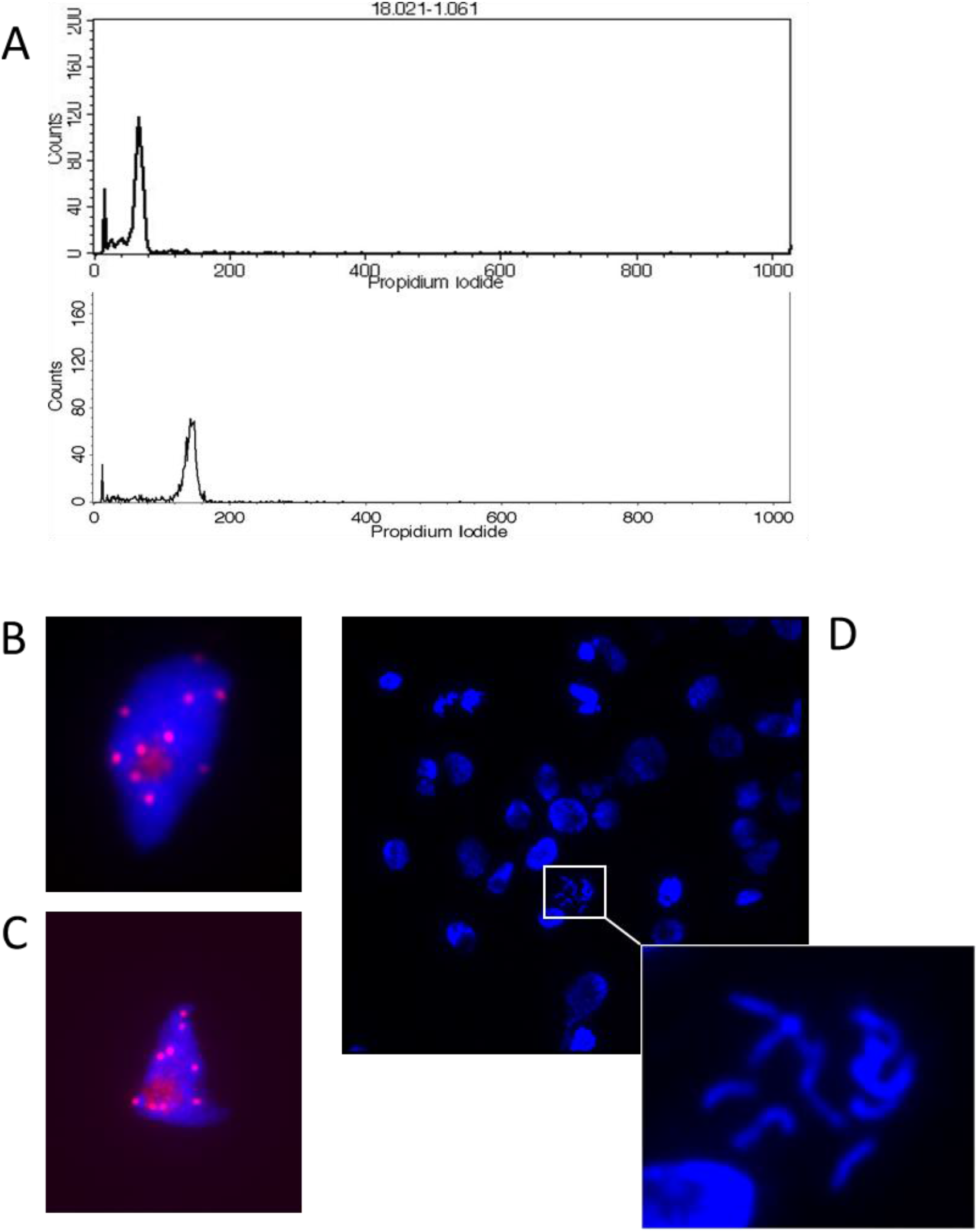
Flow cytometry (A), immunostaining with an anti-*DcCENH3* polyclonal antibody (B, C), and root tip mitosis (D) of the partially haploid plant 18.021-1

**Suppl. Figure 6.**
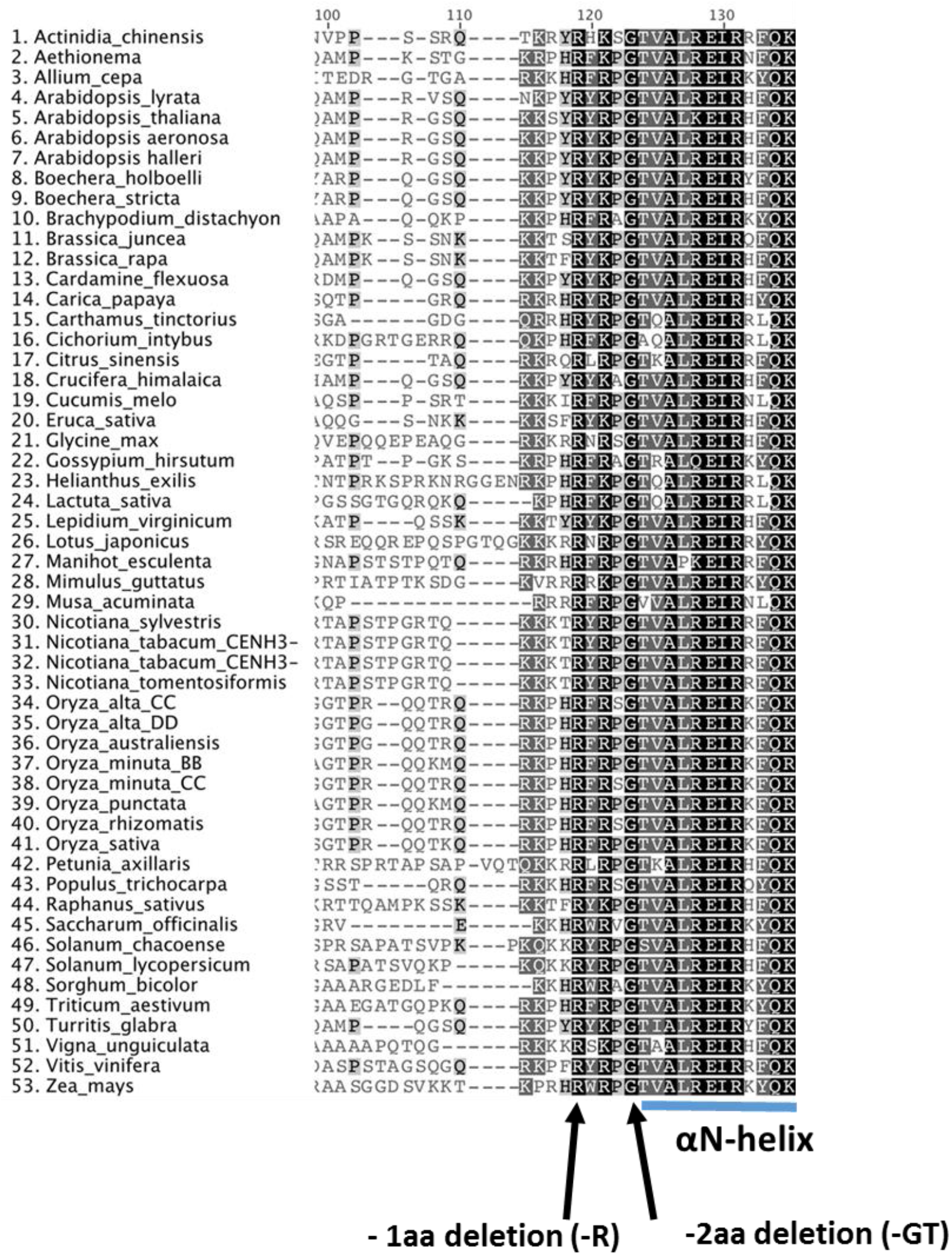
Sequence alignment of the HFD (histone fold domain) of CENH3 from 53 different angiosperm species showing the −2aa in-frame deletion resulting in haploid induction in *Arabidopsis*, and the −1 aa in-frame deletion present in carrot mutant M23-3. Sequence alignment is reproduced from Kuppu et al. (2020). The αN-helix begins at T124 (TVAL).

## Supplemental Information 1 and 2

### Suppl. Information 1

Genomic DNA sequences and coding sequences (CDS) of newly cloned carrot CENH3 genes

#### *DcCENH3A-DH1* (genomic)

ATGGCCCGAACGAAGCACCCTGCCAAACGTACCTCCGGTCACCGCTCCCGAGGTCTCTCTCCCTCACACACTTATTTTAT TAATTTAATTTGTACTGTTTTTCAATTGCTCTTTACAAGCCCTAAATTATCTTCCTCCTTTTTTGTTACAGGTCCCCCTCTCTCGGTATCTCTCTCTCTCTCTCTCTCTCCCCTCTCTTTCTCTCTCTCTCTGAAATTGGAGTTATATAAAATTTTGAATGCTGATGAGTATTTATACAGGGAACTCCGAGACGAAGAAGTACTGCTACCCCGAGTAAGTCTATAATCTTCGTATTGTTCGAGTTAGGGATTCAGTTTTTGTTAGTCAAATGAAACTGATTTCGTTGATATAGGAGAGGCAGATGCACAGGGACAACAACAGCGAAAGCCCCACAGATTTAGGCCTGGTACCGTGGCTCTTCGTGAGATTCGTAAGTTTCAGAAGACTTGGAACCTTCTAATTCCTGCTGCTCCGTTTATCCGCACTGTAAGTTATCGCTACTTGTTTAATTTCTGTTTATTTTTGTATTTGGTTTGGTATTCGGATATAATTGCCCTTAGTAATGGATGGAAATTTGAATTATATCGACTGTGGAAGTGTGAACGGATAGTAATTAAAGTTTAAATGCTTTGAACCAGAATGGCACGGGATATTGTGGTATGATTTAATTGAAGTTGTACCTAATAGGTTCTTATTTGTTGCGTACACTCAAGATCTGCTAGCTTCTTTATATAGTTAGATGCACTGATTTATTTTCATGACTAACAATGTTAATGTAGGTATTTTAAATGTTCTCAAGAAAACTTTTTATTTGGTGGATAATATGGAAAAGTTCTTTTCTTATTGAACTATTATTTGTTGAAAATTTACCCCCCAAATCTGGTTTTCTTTTCGTGTATGATAGAAGTTATTTGGCAGAATGTGTTATTATGTGAATATCAAACAACCTGAGCAACAGAATACTGATTTAATATGGACTATAATTGTAAATCTAGTTGAAATTGTCTGTAGTATCATGAGTGACTAGAATATATTGTGCATATATGTACCGCACCTTTGTAACCTACAACTCTCATTTTGAAATGGTAATAGTAATATCAAATATGTCTACTTGTGAATTCAGTTGGTATTTGATGTCATAATAATTGTTAACAGGTTAGGGAGATTAGCTTCTACCTGGCTCCCTCAATCACACGTTGGCAAGCTGAAGCACTAAGAGCAATTCAAGAGGTATTTAAAAATTCTGTCCAACTAAATTCTGTTGTTCAGTACTACTTTGTCATTTTTTTTAAAGCATTATAGCTGAATTAGAAACACTATCAAATGACCAAGACTTTAACAGTACTACTCTGGGTACCCCTTTCCGGCTTTCCTATATATAATACAAAGTTTGGCTCTGGAAGAAACACACTGCATATAGGCAAAGGCTTAATCAGCTATTTAATGTTATACTTGTCACATTTAATGCCATAAGCATACATATCAAATTTAAAAGACTTGACCAGCTTATTTTGTTTCCAAAATCATACAGAAGAAATACCATTATTTTACTCACTAGATATTGTAAATTTTCTTATGAAGCTTCCTTTGACTTATTGAAAAGATATACGATAATCCACTTAAGATAAATCGTACTCCTTAGAATTGTAAAAACAAATGATTGTGCATTTGTGCCCCTTCCTGCAATAGACGACATATTAGCTTCATGTAGAAGCCCCTTACAAATATCAAGTATTAGTACAGAATTATGCATAATGACTAAACCTTGACAATTGGAAGTTTGCAATTATTTCATTGTGGAATGAGTTTTTTGTTTCAAAAATTTATCTCTTGTGGAACAAAGTAAGTAAAATTTTATTTAATAATGAAACCAAATAACTATGTTAGTTATTTTATTTTAGAGGTTGTACAATTGACATAGATGAGCATGAGATAACAGAGCAAATAATAATTTCAAATGATATACCGCTCTTTTTTTTAAACATAACTTCTTCCATTATCGCATTCTAGGAACTGCATCTCTAACCGCATATTTTTTTGTTTAGATCAGTAAAATCGATAACCCCCGATTTTTTCGTAAACTATTTCCTTTCTCTACATGCTAGTACATAGAGTTTTCCCTTTCTTAGCTAGGCTGTTTGCATCTTTGCACTTTTAAATCTGCTCTCTAAATGTTTTTTGGAGATATTCTCTGATGTATAGAGACCTAATATCCTGGCTCACAGTTATAAATTTATTAGTCATTTTTTTGGTTATAATATATTTACAGTACCTGTGTTTGAAGTTGTTTAAATATATTTCCAGCAATGAATACCTTTTCTTTATCTGTTTTTTTATGGTTTTATCGTTGGTTATGTTTTTAGTTTTGGTCCTTCAGAGGGTAAATTGTCATGATTTTCAATAATTATAATTGTATATATTGTACTTCTCTAGATAAGAGTCGACAAAGATAACATGAAACAAAAGAAAGGTTCTTTCAAAGCGTTGATATAATCTCTGTTCTTTCAAAGTGCCATCATCATGTTTTTTAGATATGATCTCTTTTACTTATTAAGGCCCAATGTCAAATAGTTACACAGTGGCTCTGAAGCAATCTGAATTCCAGGACTCAATTTACCAATATACAAAATCTGTGCTCTACAAATCTGTTTTTGTGCTTGCAGAGTTGCAGTAATGTTCTGAGTGGAGTATCTTAGTCCTGCAAACCTTTCCACTAAACTCACGCCAACACAAGTCACTACTTACATCGTTGCATATTTAAATGTCTTATTAAGTTTTTTAGTTACTTTTTAGGTCTTTTACAAGTTAAATTCCAAAAAACACCTCCCAGACACGATAAAAGTGGTATTTCTCTTTATATCTTTAAGATTTATTACAAAAAGTGGTGTTTCTCTTGAATCCACGATTGTTCTTCCTTATTTTCTGCTGAAATTCTCACGGCTACTTGTTTGTGTTTAAATTCAATGATATAAATTGGAATCGCATATCTGAATTGCCCTAATTGTTTAGGTTCATATTGAATGATAAATTGATAACAAGTGGTGTTTTTCTTTATACCTATATGAAAATTTCTTTCAAATCCATGATTTATACTTCCTTATTCTCTGCTGAGATTCTTGCGGCTACTTGTTTGGGTTTAGATTTTATGATATTATTTGGGCTTGTATTTCTGAATTTCCTTTGTTGTTTGGCTTCATTTTACTTGAATTCTATGTGTTTAATTGAGCAGTTTATGTTGAGTGTGGTATTTTCAAGAACTGGCTCAAGAAAACCGGAATCAGAACGGATCAACTAACTTCAATTAAAGACACAGGAAAAATACATAATTCAAACGATTAGGACAAGTAAAAGATAATTCAAATAATTTCATTCGATCATATATGCATTTTAGCTGGCCAGCAGCAATTCTATACTGTGAGAATTCAGATTCTAAATTGTATGTTTTTTAGTAAAGGTACTGTTCATTTTGAAGTAGTATTATATTAATGATATTCTTTGAGTTCTCTTGAAATATTCTGGGAATCTAATATACTTGCAAAAGATATACTTTTACTTAGATGTAGCCATCTTTGCAGGCAGCAGAGGATTTTATCATTCATCTATTTGAGGATGCGATGCTATGTGCAATTCATGCAAGACGTGTCACAGTTAGTAAGTGTAAATTGATGTTTTGACATTTTTGTTTTTATCAGCCATTGCATCCCCTTTGTCTCTTTATGTTGCATCCTGAATTCATATTTAGAATATGGACTAGAGTTCGTGTTCTCTTAAATTTGCATATATATATTGAAGCTCACAGATGACAGAACCACTAGAGCATTGTACATCCTATTTCTTCTCTGTGTAGCATTTCTAAGAATTAGATTGACTTCATTAAAATTGTAAGTAATCCCAGTTAGCCATTATAGTCTCCCTGAATGGCTGCTGCACCTGTGAATTCTTGCATTACAGCTACTCACCTAGAGTTTAAATGACAAACACAATTTATAACCAGATCATGGTGACTGGCACTGGCCCAAAGCTTTGTTTAGTTGAATGAGGTATCATATACATGTAGAGATTAGCCACTACACGGGGGTTTTCTTTACTGTTTATTGTTTTTATCTGCTTTTTCATTGAGTTTGTAGAGTGTTGTAAAAAGCGGAAATCGGAATTAATCAGCGGAGCCACCGTCTAGCGATTAATAGATAAGCAGGGATATTATACTAGTTCTAAAGAAAAATGTAGTGATTAATCGGTAATAAAAAATCGAATCCGCAGAGTTCCTTGTAGAGATTAATCGGTCAAATCGGCAACTTTTAGAACTATGGTTTGTACTAGATGCAAATTGACTCAATTAAGATTTTTTTTTAGTTTTTACTAGGGTCACTATTACTCTCTGAGCAAGCAGCTACTTATTCGTTGTTATTATTGTTCTGGCTCATAGTGAAAAAAGACTGGGAGTTGGCACGCCGGCTGGGAAAGAAAGCGCAGCCCTGG

#### *DcCENH3A-DH1* (CDS)

ATGGCCCGAACGAAGCACCCTGCCAAACGTACCTCCGGTCACCGCTCCCGAGGTCCCCCTCTCTCGGGAACTCCGAGACGAAGAAGTACTGCTACCCCGAGAGAGGCAGATGCACAGGGACAACAACAGCGAAAGCCCCACAGATTTAGGCCTGGTACCGTGGCTCTTCGTGAGATTCGTAAGTTTCAGAAGACTTGGAACCTTCTAATTCCTGCTGCTCCGTTTATCCGCACTGTTAGGGAGATTAGCTTCTACCTGGCTCCCTCAATCACACGTTGGCAAGCTGAAGCACTAAGAGCAATTCAAGAGGCAGCAGAGGATTTTATCATTCATCTATTTGAGGATGCGATGCTATGTGCAATTCATGCAAGACGTGTCACAGTTATGAAAAAAGACTGGGAGTTGGCACGCCGGCTGGGAAAGAAAGCGCAGCCCTGG

#### *DcCENH3B-RO* (genomic)

ATGGCCCGAACGAAGCACCCTGCCAAACGTACCTCCGGTCACCGCTCGCGAGGTCTCTCTCCCTCACACACTTATTTTATTAATTTAATTTGTACCGTTTTTCAATTGCTCTTTCCAAGCCCTAAATTATCTTCCTCCTTTTTTGTTACAGGTCCCCCTCTCTCGGTATCTCTCTCTCTCTCTCTCTCTCTCTCTCTCTCCCCTCTCTTTCTCTCTCTCTCTGAAATTGGAGTTATATAAAATTTTGAATGCTGATGAGTATTTATACAGGGAACTCCGAGACGAAGAAGTACTGCTACCCCGAGTAAGTCTATAATCTTCGTATTGTTCGAGTTAGGGATTCAGTTTTTGTTAGTCAAATGAAACTGATTTCGTTGATATAGGAGAGGCAGATGCACAGGGACAACAACAGCGAAAGCCCCACAGATTTAGGCCTGGTACCGTGGCTCTTCGTGAGATTCGTAAGTTTCAGAAGACTTGGAACCTTCTAATTCCTGCTGCTCCGTTTATCCGCACTGTAAGTTATCTCTACTTGTTTAAATTTCTGTTTATTTTTGTATTTGGTTTGGTATTCGGATATAATTTCCCTTAGTAATGGATGAAATTTGAATTATATCGACTGTGGAAGTGTGAACTGATAGTAATTAAAGTTTAAATGCTGTGAACCAGAATGGCATGGGATATTGTGGGATGATTTAATTGAAGTTGTACCTTATCGGTTCTTATTTGTTGCGTACACTCAAGATCTCTAGCTACTTTATATAGTCAGATGCACTGATTTATTTTCATTACTAACAATGTTAATGTAGGTATTATTTTTTAAAAATTTACCCCCCAAATCTGGTTCTCTTTTCTTGTATGATACAGTAGAAGTTATTTGGCAGAGTGTGTTATTATGTGAATATCAAACAACCTGAGCAACAGAATACTGATTTAAAATGAACTACGGATTATAAATCTATTTGAATTTGTCTGTAGTATCATGAGTGACTAGAATATATTGTGCATATATACACGGCACCTTTGTAACCTACAACTCTCTTTTTGAAATGGTAATAGTAATATCAAATATGTCTACTTGTAAATTCAGTTGGTATTTGATGTCATAATAATTGTTAACAGGTTAGGGAGATTAGCTTCTACCTGGCTCCCTCAATCACACGTTGGCAAGCCGAAGCACTAAGAGCAATTCAAGAGGTATTTAAAAATTTTGTCCAACTAAATTCTGTTGTTCAGTACTACTTTGCCATTGTTTTAAAGAATTATAATTGAATTAGAAATACTATCAAATGTCCAAGAATATAACATTACTACTTTGAGTACCCCTTTCCTATATTTAATATAAAATTTGGCTCTGGAAGAAACACACTGCATATAGTTAAAGGCTTAATCAGCTATTTAATGTTATACTTGTCACATTTATGCCATAAGCATACGTATAAAATTTAAAAGACTTGACCAGCTTATTTTGTTTCCAAAATCATACAGAAGAAATAAATTATTGTACTCACTAGATATTGTAAATTTTCTTATGAAGCTTCCTTTGACTTATTGAAATGATACAATAATCCACTTAAGATAAATCATACTCCTTAGAATTGTAAAAACAAACGGTTGTGCCCCTTTCTGCAATAGACGGCATATTAGTACAGAATTATGCACAATGACTAACCCTTGACAATTGGAAGTTTGCAATTATTTCATTGTGGAATGAGTTTTTTGTTGCAAAAATTTATCTTTTGTGGATCAATGAAAGTAAAATTTTATTTAATAATGAATCCAAATAACTATGTTTAGGTATTTTGTTTTAGAGGTTGTACAATTGACATAAAGATGAGCAAGAGGATTGCATGAGATAACAGAGCAAATAATAATTTCAAATATATACCCCTTTTTTTTAAACATAACTTCTTCCATTATCGCATTCTAGGAACTGCATCTCTAACCAACAAACCACATATCTTTTTGTTCAGATCAGTTAAATCGATAACCCCCGAATTTTTCACACACTATTTCCTTTCTCTACATGCTAGTATTAGCTTTCCCTTTCTTAGCTTGGCTGTTAGCAGTTTTGCACTTTAAAATCTGCTCTCTAAATGTTTTTTGGAGATATTCTCTGATGTATAGAGACTTAATATTCTGGCTCACAGTTACAAATCTATTAGTCATTTATTTGGTTATAATATATTTACAGTACTGTGTTTTGAAATTGTTTAAAAATATTTCCAGCAATGAATACCTTTTCTTTATCTGTTTTTTATGGTTTTATTGTTTGTTATGTTTTTAGGTTTGGTCCTTCAGATGGTAAATTGTCATGATTTCCAATAATTATAATTGTAAATACTGTACTTCTCTAAATAAAGAGTCGACAAAGATAACATGAAAGTTAAAACAAAAAAAAGGTTCGTTCAAAGTATTGATATAATCTCTGTTTGGTCAAAGTGCCATCATCATGTTTTTAGATATGATCTCTTTTACTTATTAAGGCCCAATGTCAAATAGTTACACAGTGGCTCTGAAGCAATCTGAATTCCAGGACTCAATTTAATAATATACAGAATCTGTGCTCTACAAATCTGCTATTGTGCTTGCAGTAATGTTTCTGAGTGGAGTATCTTAGTCCTACAAACTTTTCCACTAAACTCACGCCAACACAAGTCACTACTTACATTGTTGCATATTTAAATGTCCTATTAAGTTTTTTTAGTTACTTTTTAGGTCTTTTACAATTTCGATTCCAAAAAACACCTCAAGAAACACGATAAAAGTGGTAGTTCTCTTTATACCTTTAGGATTTATTACAAAAAGTGGTGTTTCTCTTGAATCCACGATTGTTTCTTCCTTATTTTCTGCTGAAAATCTCACGGCTACTTGTTTGGGTTAAATTCAATGATATTAATTGGACTCGCGATCTGAATTGCCTTAATTGTTTGGGTTCATATTACTTGAATGATAAATTGATAACAAGTGGTGTTTTTCTTTATACCTATACTAAGATTACTCTTAAATCCATTATTTTTTCTTCCGTATTCTCTGCTGAGATTCTTGCGGCTACTTGTTTGGGTTTAGATTTATGATATTATTTGGGCTTGTATTTCTAAATTTCCTTTGTTGTTTAGGTTCATTTTACTTGAATTCAATGTGTTTAGTTGAGCAGTTTATATTGAGTGCGGTATTTTTAAGAACTGGCTCAAGAAAACCAGAATCAGAACAGATCAACTAACTTCAATTAAAGACACAGGAAAAATACATAATTCAAACAATTAGGACAAGTAAAAGATAATTTAAACAATTTCATTTGTTCATATATGCAATTAGCTGGCCAGCAGCAATTCTATACTGTGAGAATTTAGATTCTAAATTGTATGTTTTCTTAATAGAGGTACTGTTCATTTTGAAGTAGTATTATATTTAAAATGATATTCTGTGGGTTCTCCTGAAATAATCTGGGAATCTAATATATTTGCCAAAGATATACTATTACTTAGATGTAGCCATCTTTGCAGGCAGCAGAGGATTTTATCATTCATCTATTTGAGGATGCGATGCTATGTGCAATTCATGCAAGACGTGTCACAGTTAGTAAGTGGAAATTGATATTTTGACATTTTTGTTTTTATCAGCCATTGCATTCCATTTGTCTCTCTATGTTGCATCCTGAAGTGATATTTAGAACAGGGATTAGAGTTCGTGTTCTCTTTAATTTGCACATATATTGAAAATCACAGATAATAGAACCACTAGAGCATTGTACATTCTATTTCTTCTCTGTGTACCATTTCTAAGAATTAGATTGACTTCATTAAAATTGTAAGTAATCCCACTATCCCAGTTAGCTATTATAGTCTACCTGAATGGCTGCTGCACCTGTGAAATCTTGCATTACAATTTACAGCTACTCAGCTATAGTTTAAATTACAAACACAATTTATAACCAGATCATGGTGACTGGCACTGGCCCAAAGCTTTGTTTAGTTGAATGAGGTATCATATACATGTAGAGATCAGCCACTACACGGGGGTTTTCTTTACTGTTTATTGTTTTTATCTGCTTTTTCATTGAGTTTGTAGAGTGTTGTAAAAAACGGAAATCGGAATTAATCAGAGAGCCACCGTCTAGCGATTAATAGATAATCAGGGATTTTATACTAGTTCTAAAGAAAAATGTTGTGATTAATCGGTAATAAAAAATCGAATCCGCAGAGTTCCTTGTAGAGATTAATTGGTCAAATCGGCAACTTTTAGAACTATGGTTTATATGGTTTATACTCAATGCAAATTGACTCAATTAAGATTTGTTTTAGTTTTTACTAGGGTCGCAATTACTCTCTGAGCAAGCAGCTACTTATTCGTTGTTATTATTGTTATGGCTCATAGTGAAAAAAGACTGGGAGTTGGCACGCCGGCTGGGAAAGAAAGCGCAGCCCTGG

#### *DcCENH3B-RO* (CDS)

ATGGCCCGAACGAAGCACCCTGCCAAACGTACCTCCGGTCACCGCTCGCGAGGTCCCCCTCTCTCGGGAACTCCGAGACGAAGAAGTACTGCTACCCCGAGAGAGGCAGATGCACAGGGACAACAACAGCGAAAGCCCCACAGATTTAGGCCTGGTACCGTGGCTCTTCGTGAGATTCGTAAGTTTCAGAAGACTTGGAACCTTCTAATTCCTGCTGCTCCGTTTATCCGCACTGTTAGGGAGATTAGCTTCTACCTGGCTCCCTCAATCACACGTTGGCAAGCCGAAGCACTAAGAGCAATTCAAGAGGCAGCAGAGGATTTTATCATTCATCTATTTGAGGATGCGATGCTATGTGCAATTCATGCAAGACGTGTCACAGTTATGAAAAAAGACTGGGAGTTGGCACGCCGGCTGGGAAAGAAAGCGCAGCCCTGG

### Suppl. Information 2

Putative promotor sequences of *DcCENH3A* and *DcCENH3B*

#### DcCENH3A

ATTTGATCCAAGCCTTACGGATCACCCAAAATATCATGTAAGTCCTTAGAAACGTTAAACCCATAACAAAATACACAAAAACATCAACAAACACAAATATCTCAAAAAGATGGGTACAATCTTGATGACAAGATACTCAACAAGAATTCATTCAGAAAAAGTTAGATCTTAATTTTATTAATAAAATTGATAAATATGAGAGAAGATGAAAAAACAAAAATGATTAATTACTAAAAAAGATGAAGGTAAGAGGATGAAAGTTGAAGAAGGGATATATGGCACCTAGAGCCCGTAAGAGCAACTCCAATGCAATGCTATACTTGGTTCTATTGCTATATTATAGCATTAAAAGTAAAAAAACTCAACTCCAAAGGGGTGCTATATTTGGATGCATCCATGCATAGATGCATCCTTGGTTCTATATTTGAAACCAAAGATGGATCCATCCATGTTGCCACATCATCAATAAATAGAAATTGGAAACTAGTTAATGAGTTAATAAGTTAGGTAAAAGTTAAAGAGTTGGTTCTATATAGAAGTTAGGTAAAAGTTAATGAGTTGGTTAAATTTGAAACAAGTTATAAAACTTAGCATTGGAGTGCAAGTTTTATTTTAGCACTAAAAAACACTTTTTGGTGCTATATTATAGCAATAGCAATAGATATATAGCATCTAGCATTGGACTTGCTCTAACAGCAAATCCCGACATGGAGCAAAAATTAATCCAAGTCATGATTGAGTTTAAAATTATTAAAAATAAAAATTTATTTTGTCATCTATATTTATTTTAATCATTCGTGATTTGTAAGAACAAATATTTAATTAAGATACATATAAGATTAAATAATTTAAAAAAAGAAGAAGAATATTATGAATCCGTCCTAACAATTATATGTCGAGACATATTGTCGAAGAAAATAAACTGTTGGGTGAGTTAACTCTAAATGCAAGATAATCCAGTAACAAAAATGACATTTTAGGATTTTAATTACGAACCACTGAATTTATAAACAGTACACGTAGACTGAAAATTAGGACAGCGGGAACGAAAATTTGGTCAGTGGCAGTAACGGCTCTTTCGACGTTTAAATCACTCCCTCCGTCTTATTTCAAATTAATCGACAACCATTTGGAATAAAAAAAAATCATAATTTTATAACCGCCCGTTAGAAATCACGGTCATCATTTCCGGCAAGTG

#### DcCENH3B

TATAGCACTTTTCCCCCAGTTATGCCAGTCCTATTGGAGTGTGCTATATATAATCCTTTACAACGTGTAATGTGTACTTGTTTAATAGTAGCAAAATTATGAAATTAATATAATTTAACCAATGTTTCATTATATTTAGTTTCTTTTGGAACTTTTGTGGTTATCAGAATGTTGCTTATCTTGGTAAAAAAATAGCTAAGTATTTAATCCGGAATTAATTTTACTTGATACTATCTTTTACTGAAATTCATAGAACATTTTCTAAAAAGTTAAATAAACATATATTAAATCTTGGTACAAGTATCTTGTAGGATTCAACAAAACATGAATAGTTAAAAAGTTAAATTAAATAAAATATTTTTCGGAAAAACTTTTTCGCACTAACGAGCTTTGACATCTATACTATAATATAATAAGTCAACATGAGTATAATTTGTAGTTCAAGATTTTAGTTATACTAACCTTTAACCCGTGCGAAGCACGGGTGGGTATATAGTTCGTAATTTAACATTATTTTATTGGTATTTTAGTATTAATGGATTGAATTTTAATTAAATTATATTATTCATCTGACTAGATTACCGAATCTTTCAAAGGTATTACACGATCGGCTATGTTTGGAAAAGATTAGAATTTTCTGATTTTTTTTATTTTTAAATTTATGTGTACTAAATTCGGAATATATATATATATATATATATATATATATATATATATATATATATTAGGGGTTGTCCAATTTTAAGTAATCGGGAGAAAATATAGTCAGTTGAATAATATAATTTAATTAAAATTCAATCCATTAATATTAAGATACTAATAAAATGATGTTAAAATTCAAATTATAAATAAGAAATTACGAACTATATACCCGCCCGTGTTTCGCACGGGTTAAAAGGCTAGTCCAATCTTAACCATAGAAATGAGCCTATCTCAAGTTTTCATTTTAAAGTACTGTTTCTGTTTTTTTTTTTTTTCGGTTTTTATTTTATTTTAATCTATCTACTATATCTATTATACGAGAACAGGTTAGTTTACTGAGCAAATTAATTCCCGCTTCGGTCGTTCCAACCCTCCTTCCAAACCTGCTCATTAAACCCCATGTATTACCCATCTACATACAAACAGACATCTCCCTTCAGTTGAAATCCACGGTCATCATTTCCGGCAAGTG

